# Limitations of *de novo* sequencing in resolving sequence ambiguity

**DOI:** 10.1101/2025.08.19.671052

**Authors:** Sam van Puyenbroeck, Denis Beslic, Tomi Suomi, Tanja Holstein, Thilo Muth, Laura L. Elo, Lennart Martens, Robbin Bouwmeester, Tim Van Den Bossche, Tine Claeys

## Abstract

De novo peptide sequencing enables peptide identification from fragmentation spectra without relying on sequence databases. However, incomplete spectra create ambiguity, making unambiguous identification challenging. Recent deep learning advances have produced numerous de novo models that predict sequences and refine peptide–spectrum matches under such conditions. Yet, their relative strengths, weaknesses, and ability to handle spectrum ambiguity remain unclear. Here, we benchmark eight state-of-the-art models on three publicly available proteomics datasets, comparing performance using established metrics and quantifying inter-model agreement. We assess post-processing approaches, including iterative refinement, rescoring, and reranking, for their ability to improve identification accuracy, and perform an error analysis to identify common mispredictions and their causes. Model performance varied, with considerable overlap of correct identifications. Post-processing yielded no or only modest improvements. Most sequencing errors were model-independent and driven by limited fragment ion coverage, a limitation also observed in database searches with large search spaces.

## Introduction

High throughput proteomics heavily relies on mass spectrometry to detect and quantify peptides, which are then used to infer protein presence and abundance^1,2^. This process begins by identifying peptides in acquired fragmentation (MS2) spectra, obtained by fragmenting precursor peptides that enter the mass spectrometer. Each peak in an MS2 spectrum has an associated mass-to-charge ratio (m/z) and intensity value, most of which are related to the amino acid sequence of the fragmented precursor peptide. A peptide search engine then uses the predictable fragmentation behavior of peptides to score peptide spectrum matches (PSMs) for the recorded MS2 spectra^3–7^. However, even with state-of-the-art instruments and identification algorithms, the exact peptide sequence cannot always be unambiguously determined^8^.

Three main factors contribute to this challenge of identification ambiguity. First, some peptides generate b-ions and y-ions of (nearly) identical mass and are thus notoriously difficult to distinguish by mass spectrometry. Well-known examples include isobaric amino acids such as leucine and isoleucine, as well as pairs that become isobaric through post-translational modifications (PTMs), such as aspartic acid and deamidated asparagine, or glutamic acid and deamidated glutamine. Nearly isobaric pairs such as oxidized methionine and phenylalanine present similar challenges. Resolving these cases requires additional spectral evidence beyond b-ion and y-ion m/z values. Secondly, sequence ambiguity increases when ion ladders, i.e. series of consecutive fragment ions, are missing. In particular, when no fragment ion is observed at a specific cleavage site in either b-ion and y-ion series, i.e. a missing complementary ion pair, the site remains unannotated. The absence of such complementary ion pairs significantly increases the number of possible isobaric sequences that can explain the observed spectrum well^8^. This often results from suboptimal fragmentation, influenced by, amongst others, collisional energy settings or precursor ion properties. Thirdly, the increase of fragment peaks due to co-fragmentation or experimental noise in the spectrum can be detrimental for unambiguous identification. New, yet spurious amino acid ladders can appear, increasing the number of likely peptide candidates.

The most common approach to mitigate sequence ambiguity is to dramatically limit the search space to only the very small subset of peptide sequences that are expected in the sample^9^. This is achieved through sequence database searching, where observed spectra are matched only against sequences from the predefined protein database^10–14^. Although an effective way to reduce the search space and therefore minimize sequence ambiguity, sequence database searching also brings several limitations.

One key constraint of sequence database searching is the inability to identify proteins or isoforms that are not part of the database. In proteogenomics, this issue is addressed by including additional sequence variants derived from genomic and transcriptomic data in the search database^15^. Similarly, metaproteomics workflows often rely on large, across-species sequence catalogs, or even use entire reference databases such as UniProtKB or NCBI RefSeq^16^. However, this process of expanding the database directly increases the potential for identification ambiguity by again increasing the number of distinct candidate peptide sequence matches^17,18^. Indeed, growing the sequence repertoire of the search database makes it a less efficient filter for resolving potential sequence ambiguity. Interestingly, recent machine learning-based rescoring approaches such as MS²Rescore have shown great promise in resolving identification ambiguity in searches against large databases^19^. These approaches achieve this by not only considering b-and y-ion mass-over-charge (m/z) information, but also fragment intensity patterns^20,21^ alongside other matching information such as the peptide precursor’s retention time^22^ and collisional cross section (CCS)^23–25^. At their core, these rescoring approaches compare the experimentally observed values with predictions for these values made by machine or deep learning models. By integrating these features into a scoring model trained to separate likely true from known false, i.e. decoy matches, more reliable identifications can be achieved even in complex databases^26,27^. Even though these approaches work well to maintain identification performance in large databases, they remain reliant on databases and are thus unable to identify novel peptides or unexpected variants.

*De novo* sequencing is a promising alternative, as it does not rely on sequence databases and can thus, in theory, identify any peptide from a fragmentation spectrum. However, because *de novo* sequencing essentially removes all constraints on the search space, the ambiguity issue returns in full force. Peptide sequence generation, feature extraction and PSM scoring must therefore be handled with care to ensure accuracy. Recent deep learning-based models have pushed *de novo* performance to new levels^28^. These state-of-the-art neural networks predict peptide sequences in an autoregressive manner, generating one amino acid at a time based on the previously predicted ones. They use beam search to explore multiple candidate sequences in parallel and apply precursor mass control to ensure consistency with the observed mass. Since the development of the Casanovo model^29^, the application of attention mechanisms to *de novo* peptide sequencing has become central to learn the complex relationships between the intensities and masses of peaks in an MS2-spectrum and the peptide subsequence under construction. Nevertheless, questions remain regarding (i) what the strengths and weaknesses of different *de novo* models are, and (ii) to what extent they can deal with the identification ambiguity problem.

In this study, we therefore systematically benchmark eight deep learning-based *de novo* sequencing models (AdaNovo^30^, Casanovo^29^, ContraNovo^31^, InstaNovo^32^, InstaNovo+^32^, NovoB^33^, PepNet^34^, π-HelixNovo^35^, π-PrimeNovo^36^) alongside three post-processing methods (InstaNovo+^32^, MS²Rescore^19^, and Spectralis^37^). First, we evaluate their ability to reproduce database search results. We then assess the extent to which post-processing methods, including rescoring and reranking algorithms, can improve the original *de novo* results. In cases where de novo and database-derived PSMs disagree, we analyze how prediction errors relate to specific spectral characteristics. Finally, we demonstrate that although *de novo* sequencing excels under specific conditions, its limitations, particularly with regards to low-quality spectra, mirror those of database-based approaches.

## Results

Recent advances in deep learning have led to a surge of new deep learning-based *de novo* models. To systematically evaluate their performance, we structured our analysis into three sections. First, we benchmarked eight state-of-the-art models using commonly used evaluation metrics and assessed their agreement across three publicly available proteomics datasets.

Second, the extent to which post-processing methods can improve the initial *de novo* results are assessed. These include models that iteratively refine candidate PSMs, rescore PSMs, and rerank candidate sequences. Each approach is benchmarked in terms of their ability to increase identification accuracy and improve the ranking of the correct PSMs.

Finally, a detailed error analysis to characterize the limitations of current models is performed. Here, the most frequently occurring prediction errors for each model are reported and associated with spectral features. This will provide insights into specific spectral features under which *de novo* sequencing remains challenging.

## 1 Assessment of accuracy and consensus across *de novo* models

In *de novo* sequencing, evaluation metrics such as peptide and amino acid accuracy, recall, and coverage are commonly used when comparing the performance of *de novo* models^28^. Here, we assess coverage based on the spectra identified with a database search, unless otherwise specified. Alternatively, coverage can be defined as spectra identified by both database search and all *de novo* models. This distinction is important as not all models generate predictions for every spectrum, some predictions are filtered *post hoc* because of precursor mass mismatch, and some PSMs are simply not considered by the *de novo* model due to unsupported modifications. In this study, we use identifications from a Sage^12^ search against a combined *Homo sapiens*, *Escherichia coli*, and *Saccharomyces cerevisiae* database, (or, in the case of the metaproteomics dataset, a FASTA provided by the authors who generated the data), filtered at q-value < 0.01, as our gold standard.

Beyond overall accuracy, we also evaluate *de novo* model complementarity by measuring the overlap and uniqueness of the correct predictions made by each model. Measuring agreement and disagreement identifies (dis)advantages of model architectures and indicates whether the combination of different models in an ensemble could improve results.

Three datasets are used for benchmarking. Two datasets originate from a quantification benchmark consisting of a *Homo sapiens*, *Escherichia coli*, and *Saccharomyces cerevisiae* mixture (PXD028735), acquired on an Orbitrap QE-HFX and on a timsTOF Pro respectively^38^. These two instruments represent two distinct fragmentation and mass analysis technologies, which substantially change the acquired data. The third dataset contains samples spiked in with a mock community of eight species (PXD023217) from the CAMPI study, analyzed on the Orbitrap QE-HF and Fusion Lumos. This dataset simulates a metaproteomics experiment using a mock community composed of defined species.

### 1.1 Accuracy

*De novo* model performance is evaluated based on how well correct peptide predictions are prioritized, i.e. scored higher, over incorrect or low-confidence predictions across spectra. This performance can then be visualized on a precision-coverage (PC) curve, which shows the precision at several score thresholds. The coverage indicates how many spectra acquired PSMs with higher scores at a given score threshold. The PC curves show that PepNet consistently underperforms (Figure 1). The other models show similar performance and largely follow a consistent ranking, from lowest to highest on the PXD028735-Orbitrap dataset: PepNet – ContraNovo – π-HelixNovo – AdaNovo – NovoB – Casanovo – InstaNovo ∼ π-PrimeNovo. It should be noted, however, that on the other two datasets the precision of InstaNovo is much lower than that of the directly competing tools Casanovo and π-PrimeNovo.

**Figure 1.**
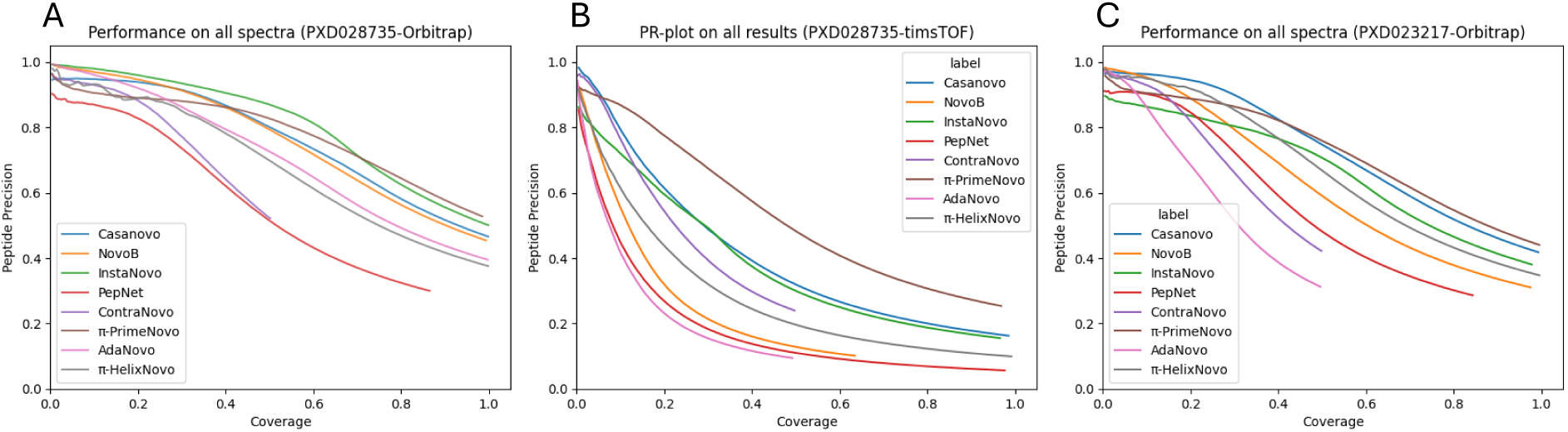
PC-curves of all eighth de novo models for three datasets: (A) PXD028735-Orbitrap, (B) PXD028735-timsTOF, and (C) PXD023127-Orbitrap. PSMs in gold standard: 186101 (A), (B) 312311, and (C) 310560; Number of samples: (A) 4, (B) 4, and (C) 6.

Interestingly, π-PrimeNovo shows an enrichment for high-scoring but incorrect PSMs, indicating overconfidence in some of its predictions. ContraNovo shows a high accuracy in the PC-curve relative to the other models, but this is not unexpected: built-in processing reduces the number of considered spectra in ContraNovo to roughly half the number considered by the other models, which leads to its PC-curve stopping at 0.5 coverage.

Precision across all models is strongly affected by the instrument and sample complexity. A noticeable drop in performance is observed for all models on the timsTOF data, likely reflecting model training biases toward HCD-based data such as the nine-species benchmark^39^ or the MassiveKB collection of HCD-spectra. This highlights limitations in model generalization to alternative fragmentation methods for all models. Additionally, InstaNovo and AdaNovo show a remarkable drop in performance in a complex species sample compared to the *Homo sapiens – E. coli – S. cerevisiae* mix, highlighting potential overfitting on species-specific sequence information.

Based on the lower overall performance across models on the PXD028735-timsTOF and PXD023217-Orbitrap datasets, we will focus on comparing model performance on the PXD028735-Orbitrap data. This ensures that all models are evaluated in optimal conditions.

### 1.2 Overlap

While all models show similar overall performance, they can diverge substantially in the specific spectra they correctly sequence. In total, 67.9% of all spectra are correctly predicted by at least one model. As shown in Figure 2A, there is good overlap in a subset of correct predictions: 12.9% of all correctly identified spectra are correctly sequenced by all models. After excluding ContraNovo and PepNet due to incomplete output and lower performance respectively, this proportion increases to 34.7%. This high degree of shared identifications suggests that there is a subset of spectra that are straightforward to sequence. Each model also correctly predicts a unique set of spectra (Figure 2B). For instance, π-PrimeNovo, NovoB, and InstaNovo each contribute 3.66%, 3.25%, and 3.15% unique correct identifications, respectively.

**Figure 2.**
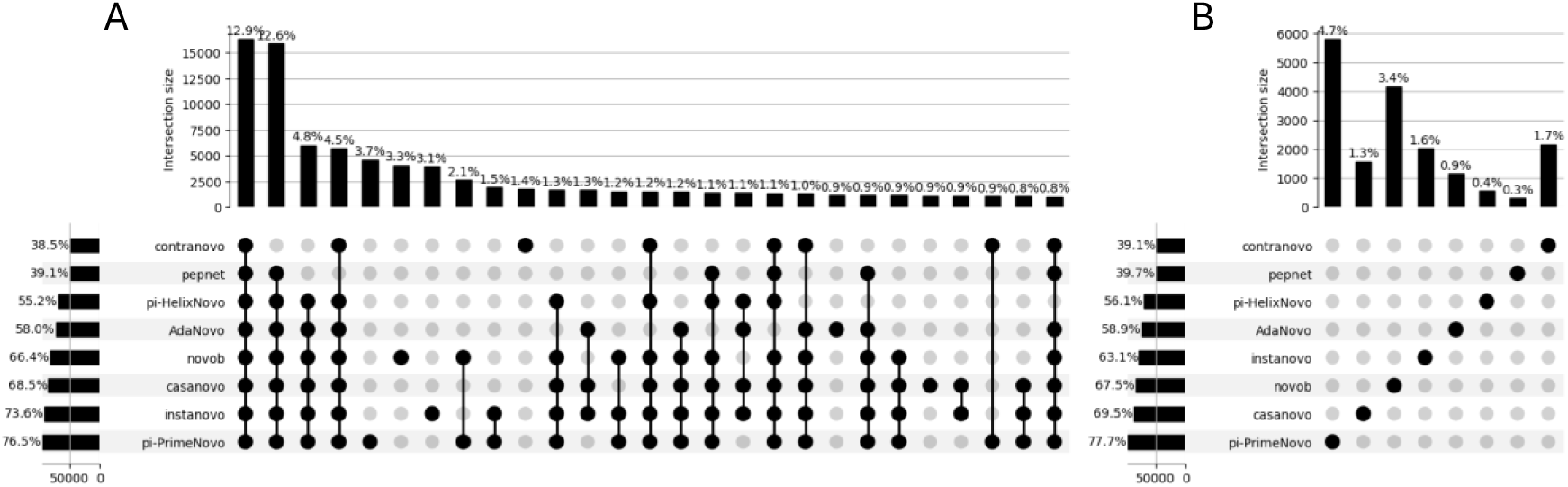
(A) Overlap of correct predictions ordered by highest cardinality. (B) Set of unique, correctly predicted spectra for each model.

To evaluate agreement between models, the intersection of correct model predictions was calculated between each pair of models. This intersection was normalized by the total number of correct predictions for each model, resulting in two values: (i) The proportion of correct predictions of model A also identified by model B, and (ii) vice versa for model B (Figure 3).

**Figure 3.**
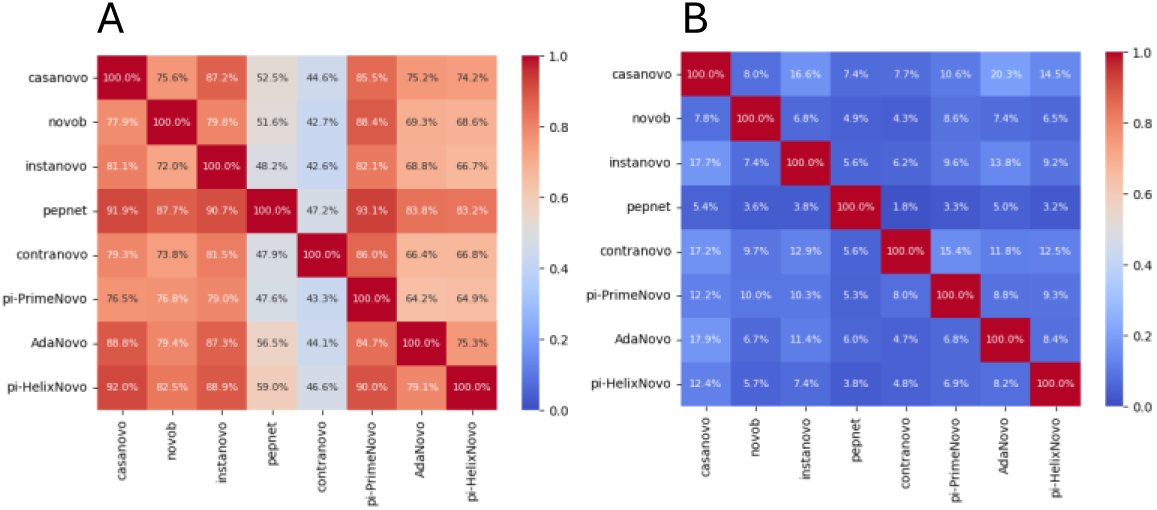
Pairwise overlap of spectra acquiring (A) correct, and (B) incorrect de novo PSMs. Each value represents the proportion of predictions from the model on the vertical axis, which were also found by the model on the horizontal axis ([N_prediction(vertical)_ ∩N_prediction(horizontal)_] / N_prediction(vertical)_).

Π-PrimeNovo, the most accurate model, encompasses at least 82.1% of the correct predictions made by any other model (vertical axis). Casanovo similarly captures ≥92% of correct predictions from both π-HelixNovo and PepNet, making the latter two largely redundant. In contrast, PepNet and ContraNovo show lower intersections, recovering at most 59.0% and 47.2% of correct predictions from other models respectively. For ContraNovo, this is partly attributable to its reduced number of reported predictions.

Intersection analysis on incorrect predictions reveals minimal overlap, suggesting little agreement when models make errors. Nonetheless, supplementary Figure S1 highlights a few cases where incorrect predictions align across all models, but this only occurs for 0.375% of spectra. These cases suggest the presence of co-fragmenting peptides, as 81.2% of these *de novo* peptides are also present in the FASTA database.

## 2 *De novo* post-processing tools

Recently, refinement models that build on top of *de novo* predictions have gained attention^32,37,40^. These models take as input both the experimental spectrum and the *de novo* output, i.e., a candidate PSM. Most post-processing approaches operate in two distinct yet tightly linked ways: (i) PSM rescoring, and (ii) candidate selection.

In PSM rescoring, only the highest-scoring candidate per spectrum is considered. A good PSM rescoring method will achieve a higher proportion of true hits at any given score threshold, reflected by a larger area under the PC-curve. This approach, already widely used for database search results, has been shown to typically improve confident identifications by 20-35% in large database searches^41,42^.

In candidate selection, the goal is to select the best PSM for each spectrum. This can be done in two ways. The first requires that a *de novo* search engine proposes a large number of candidates, from which the best one is then selected in the refinement, based on a score that describes the quality of the match. Several post-processors, including pNovo3^43^ and PostNovo^44^, have been developed to assign a new PSM score, with the objective to improve the ranking of the proposed candidates. Alternatively, some models, such as Spectralis and InstaNovo+, do not rely on a predefined large list of candidate peptides and can instead start with an initial candidate as a guide to search autonomously for a better one through iterative candidate refinement.

We will here benchmark the application of Spectralis, InstaNovo+, and MS2Rescore on all *de novo* results for both PSM rescoring and candidate selection on the extent to which they can improve the original *de novo* results. Spectralis and InstaNovo+ were used to rescore and refine the sequence of the PSMs. Both the original *de novo* PSMs from all models and the refined PSMs were rescored using an adapted MS2Rescore-pipeline, which is originally developed for refinement of database searches (see Methods for details).

Note that we refer to the gold standard, database search-derived PSMs as *PSM_Gold-standard_* and to the PSMs scored highest by any other scoring method as PSM_<scoring method>_.

### 2.1 Sequence refinement

In this section, the ability of Spectralis and InstaNovo+ to refine candidate *de novo* PSMs towards *PSM_Gold-standard_* is evaluated. Ideally, refinement models should only refine incorrect candidate PSMs, while correct candidate PSMs should remain unrefined. In addition, when refinement occurs, regardless of whether this leads to *PSM_Gold-standard_*, it may indicate that the spectrum is ambiguous, as multiple PSMs could match the observed spectrum well.

Across the eight evaluated *de novo* models, a substantial fraction of PSMs are refined by Spectralis (27.3-43.4%) and to a lesser extent by InstaNovo+ (7.62-37.7%). InstaNovo+ was notably more effective than Spectralis in identifying candidate PSMs that did not require refinement, as evidenced by the lower proportion of correct yet refined PSMs. Additionally, for *de novo* PSMs produced by PepNet, Spectralis left 54.7% incorrect candidates unrefined, compared to 48.9% for InstaNovo+, indicating InstaNovo+ is slightly more selective for refining *de novo* PSMs. Similar trends were observed for PSMs produced by other *de novo* models. In Figure 4A and Figure 4B, the effect of refinement with Spectralis and InstaNovo+ is visualized for *de novo* results generated by Casanovo. Similar figures for the other models can be found in Supplementary Figure S2

**Figure 4.**
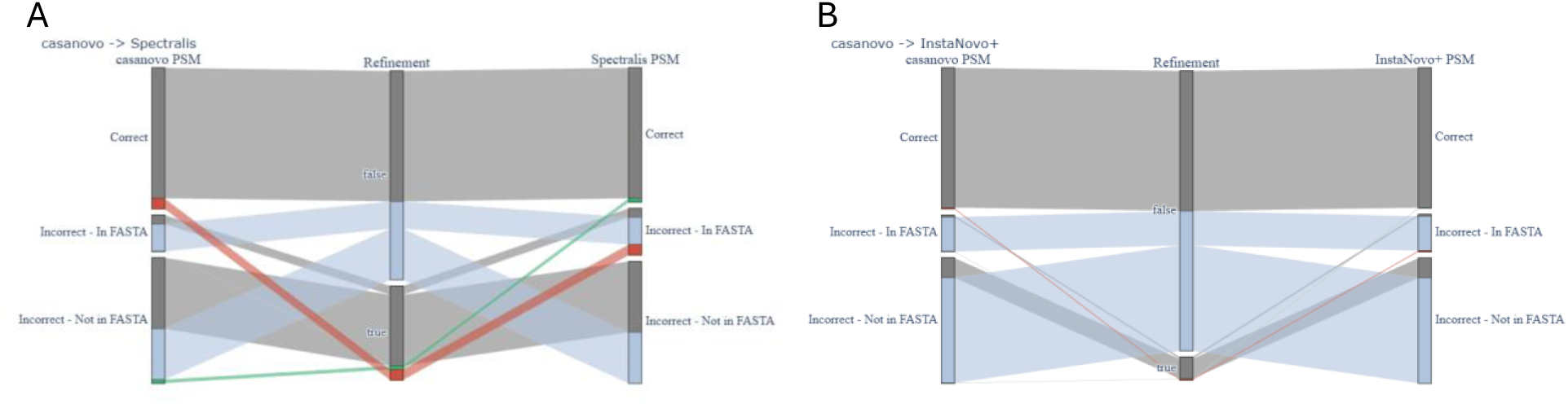
Sankey plots showing the effect of refinement for (A) Spectralis, and (B) InstaNovo+ on PSM-correctness for de novo PSMs generated by Casanovo. Red indicates changing a correct prediction to an incorrect one. Green indicates correct refinement Grey indicates a status quo situation, i.e., correct left unrefined and incorrect prediction refined to a different incorrect sequence. Blue indicates wrong predictions which the post-processor did not try to refine.

Despite frequent refinement, the increase in the number of correct *de novo* PSMs after refinement is limited. Depending on the accuracy of the base *de* novo model, both Spectralis and InstaNovo+ often yielded only minor improvements or even reduced overall performance. Particularly for more accurate *de novo* models such as π-PrimeNovo and Casanovo, refinement decreased the number of correct PSMs. Interestingly, this decline in performance was due to a combination of correct refinements of incorrect PSMs and incorrect refinements of initially correct PSMs. Importantly, these incorrectly refined PSMs were always peptide sequences which were present in the FASTA database, highlighting the issue of identification ambiguity. In general, for all models, these results indicate that refinement models are not capable of meaningfully improving upon the *de novo* prediction models themselves.

To better understand the refinement behavior, we categorized refinements into four scenarios: (i) incorrect PSMs left unrefined, (ii) correct PSMs incorrectly refined, (iii) incorrect PSMs correctly refined, and (iv) incorrect PSMs incorrectly refined (Figure 5). For each scenario, the Levenshtein distance was calculated between the refined PSM and the gold standard identification to assess the refinement steps still required to reach the solution. To evaluate whether the models were expected to find the correct solution, we computed the difference in PSM-score as defined by MS2Rescore between the gold standard PSM and the *de novo* PSM. A larger positive score indicates MS2Rescore is able to identify the gold standard as a better match (See Methods for details on MS2Rescore-rescoring).

**Figure 5:**
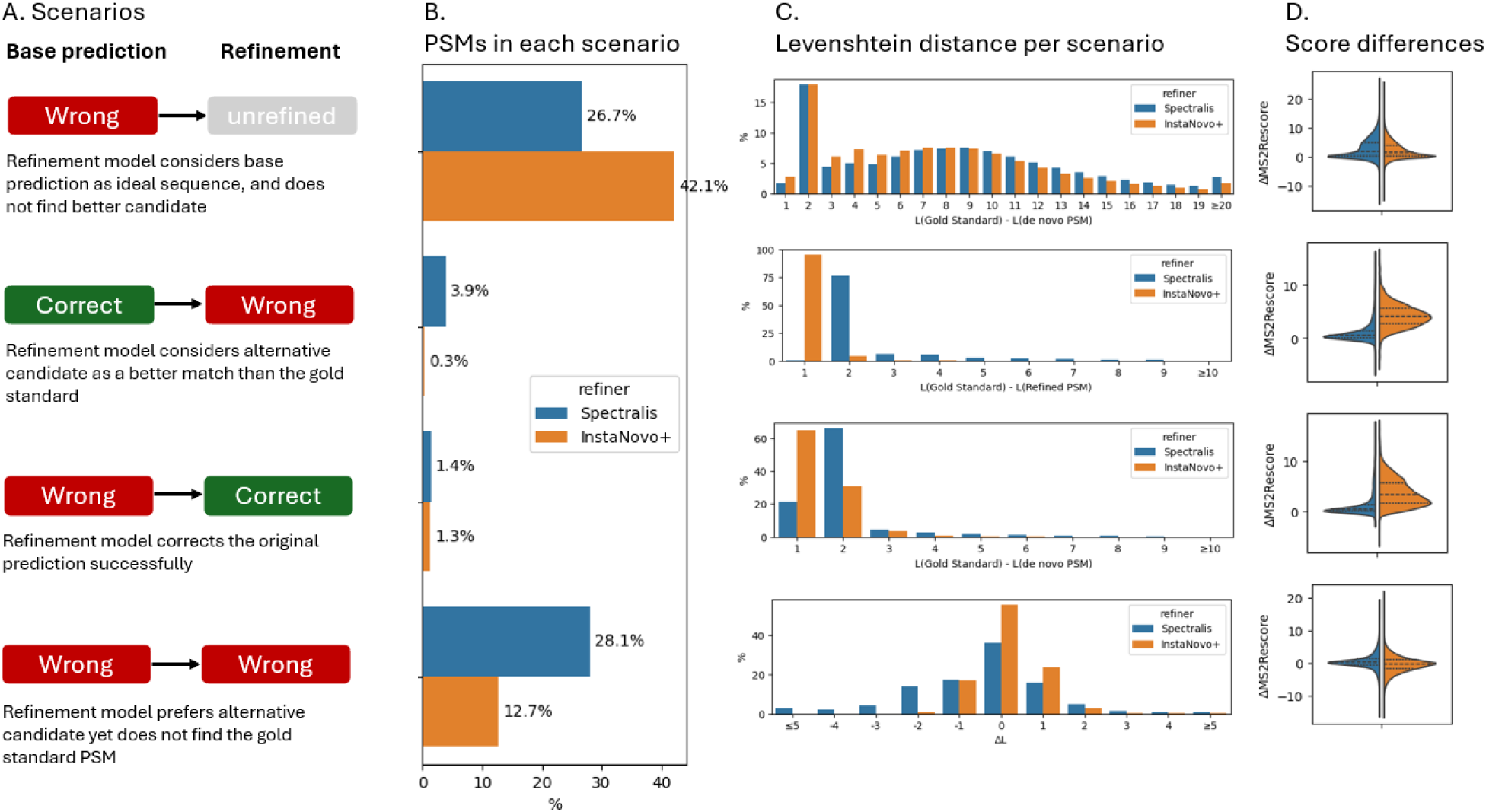
Average effect over the eight de novo models after applying iterative refinement with Spectralis and InstaNovo+. Refinements are subdivided into four distinct categories (A), and occur in unequal quantities (B). C) Levenshtein distance between gold standard and de novo or refined PSM for each scenario. In the final scenario (wrong → wrong), difference between Levenshtein distance between the base de novo and refined PSM with gold standard is shown. D) Shows MS2Rescore-score difference, computed similarly to Levenshtein distance in (C).

In the first scenario, when post-processors leave incorrect PSMs unrefined, *de novo* PSMs have a Levenshtein distance to the gold standard identification of two or less in approximately 19.5% and 20.7% of cases for Spectralis and InstaNovo+ respectively, indicating that the *de novo* PSMs deviated by no more than two amino acids. Additionally, PSMs left unrefined by Spectralis show significantly larger MS2Rescore-score differences than InstaNovo+. These results indicate that InstaNovo+, more than Spectralis, leaves PSMs unrefined when MS2Rescore too has more difficulties indicating the gold standard PSM as a better match.

In the second scenario, both Spectralis (3.93%) and InstaNovo+ (0.33%) occasionally incorrectly refine initially correct PSMs. In 77.2% and 99.8% of these cases for Spectralis and InstaNovo+ respectively, these correct PSMs are changed by one or two amino acids, even though MS2Rescore recognized the gold standard PSM as a better match, especially for the InstaNovo+-refined PSMs.

Ideally, as defined in the third scenario, all incorrect *de novo* PSMs would be correctly refined. However, this occurs in only 1.36% and 1.33% of all incorrect PSMs for Spectralis and InstaNovo+, respectively. In 92.0% (Spectralis) and 99.2% (InstaNovo+) of these cases, the *de novo* PSMs had a Levenshtein distance of three or less. This indicates that the evaluated refinement models are only able to correct very minor sequencing errors.

Finally, in 28.1% (Spectralis) and 12.7% (InstaNovo+) of incorrect PSMs, PSMs are refined, yet remain incorrect. On average, the refinement does not reduce the Levenshtein distance towards the ground-truth. Furthermore, MS2Rescore also does not consistently score either the base *de novo* PSM or the refined PSM higher. This indicates that refinement is neither able to reduce the sequence error of an erroneous PSM candidate, nor able to find a sequence matching the experimental data better based on relevant MS2Rescore features.

In summary, the performance gains achieved by iterative candidate refinements are marginal to non-existent, depending on the performance of the base *de novo* model (Supplementary Figure S3). When gains are observed, these are mostly limited to small corrections involving fewer than two amino acid substitutions.

### 2.2 Rescoring

While sequence refinement provided no remarkable improvements, we next evaluated the effect of using the refinement models and MS2Rescore as rescoring methods applied to the *de novo* results both with and without sequence refinement. An effective rescoring method should assign higher scores to correct *de novo* PSMs and lower scores to incorrect and potentially spurious PSMs resulting from spectrum ambiguity. Although rescoring does not alter the sequence itself, and therefore does not change the overall accuracy at 100% coverage, it may substantially improve the ranking quality of the PSMs, which makes setting score thresholds for result filtering more effective. Improvements made by rescoring are reflected in the area under the precision-recall curve (AUC), where a higher AUC indicates better discrimination between correct and incorrect PSMs. To evaluate rescoring performance, Spectralis, InstaNovo+ and MS2Rescore were used to assign new scores to the top candidate PSMs generated by each *de novo* model, both before and after iterative candidate refinement (Figure 6). For InstaNovo+, improvements are always a combination of iterative candidate refinement and rescoring, as the model does not support the use of both functionalities in isolation.

**Figure 6:**
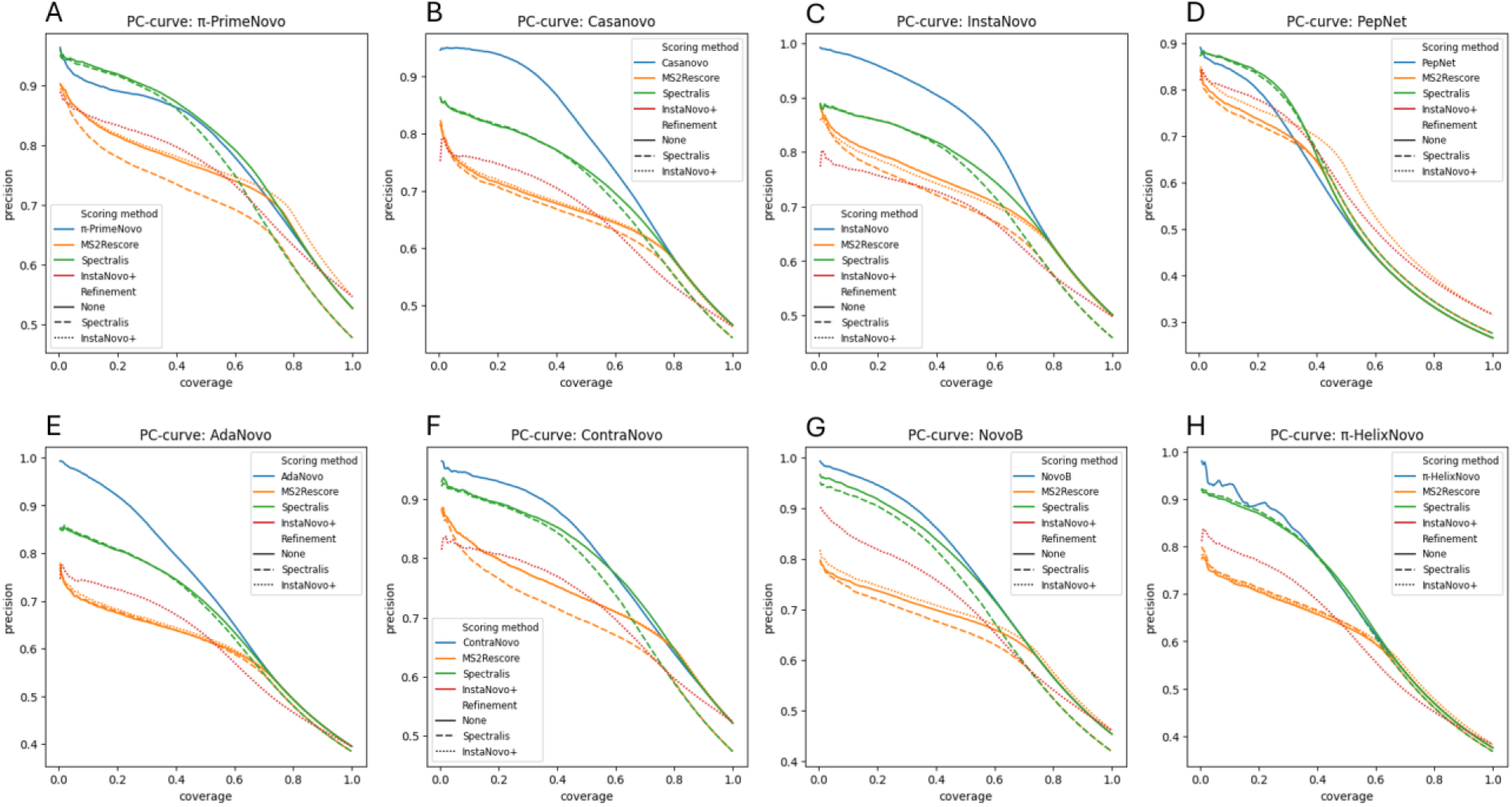
Precision-coverage curves for (A) π-PrimeNovo, (B) Casanovo, (C) InstaNovo, (D) PepNet, (E) AdaNovo, (F) ContraNovo, (G) NovoB, and (F) π-HelixNovo showing the precision coverage for four scoring methods and two refinement models.

For all models except PepNet, the scores assigned by Spectralis were less effective in separating correct from incorrect predictions. This suggests that the Levenshtein distance estimator used by Spectralis does not outperform the native scoring functions of most *de novo* models. In contrast, InstaNovo+ rescoring achieved higher AUCs for all models except Casanovo. Because InstaNovo+ could not be used as a rescoring method as such, these improvements can be attributed to both PSM refinement and rescoring. However, as the previous section only showed limited benefits from iterative candidate refinement, most improvements are likely attributable to its rescoring capabilities. Finally, MS2Rescore consistently underperformed across all models and refinement settings. It failed to better distinguish correct from incorrect PSMs than the base model scores or those from Spectralis and InstaNovo+. This is notable, given that MS2Rescore is widely used to assess target-decoy separation in database search results, a highly similar task. One possible explanation for this is that incorrect *de novo* PSMs often show considerably better matches than decoy PSMs, and are therefore not as easily distinguished as in traditional database searches. This highlights a limitation of applying a target-decoy driven method such as MS2Rescore in a *de novo* context.

### 2.3 Candidate selection

*De novo* sequencing models aim to select the best matching PSM for each spectrum from a vast space of possible peptide sequences. To make this tractable, these models rely on search strategies, filters, and heuristic scoring. Autoregressive models, such as Casanovo, use beam search to keep the top k scoring hypothesis, or ‘beams’, at each step in the peptide sequence generation process, where each beam’s score consists of an aggregation of amino-acid level probability scores. In contrast, non-autoregressive models such as π-PrimeNovo and PepNet instead decode the peptide sequences from a full amino acid probability matrix using custom algorithms. These often combine precursor mass error minimization alongside scoring heuristics. In both model types, limitations in the search heuristic or scoring accuracy can result in the correct PSM either being excluded from the candidate list or ranked below incorrect alternatives.

To systematically assess these limitations, we used Casanovo to generate ten candidates per spectrum and, where applicable, assigned a Casanovo score to the gold standard PSM when this PSM was absent from the candidate list. Spectra lacking the gold standard PSM in the original top ten candidates are denoted as *Spec_absent_*; the remainder as *Spec_top10_*. Each candidate list was then rescored and re-ranked with MS2Rescore, Spectralis, and InstaNovo+. We evaluated two feature sets for MS2Rescore: the default model (MS2Rescore-d) and an extended model that also includes the Casanovo and Spectralis scores (MS2Rescore-CS). Finally, to understand why some methods succeed or fail, we correlated ranking outcomes with spectrum-level features (e.g. fragment ion coverage, noise level) and peptide-level differences (e.g. Levenshtein distance, isobaric variants). In this analysis, we allowed *de novo* PSMs which differed only by an isobaric amino acid (such as aspartate and deamidated asparagine) to be considered as a match.

Casanovo did not report the gold standard PSM amongst its top ten candidates for 41.6% of spectra. However, when the gold standard PSM was included, Casanovo and MS2Rescore-CS were the most performant in ranking the gold standard as the first rank (Figure 7A). Casanovo correctly ranked 79.7% of *Spec_top10_*, while InstaNovo+ only correctly ranked 74.5%, MS2Rescore-d 70.0%, and Spectralis 59.6%. In contrast, for *Spec_absent_*, MS2Rescore-CS and MS2Rescore-d correctly ranked the gold standard PSM as a first rank for 75.8% and 67.2% of *Spec_absent_* respectively, outperforming Casanovo (42.0%) (Figure 7B). Notably, Casanovo flags predictions not matching the experimental mass of the precursor peptide by summing the score with-1 as a post-processing step. After removing this processing-step, which effectively works as a precursor mass error filter, 96.4% of *Spec_absent_* for which the gold standard PSM were initially ranked as the highest scoring candidate by Casanovo, acquired a different top candidate. Overall, the MS2Rescore-CS approach outperformed any other method for candidate selection, closely followed by MS2Rescore-d. Spectralis and InstaNovo+ did not improve upon the original candidate selection results provided by Casanovo.

**Figure 7:**
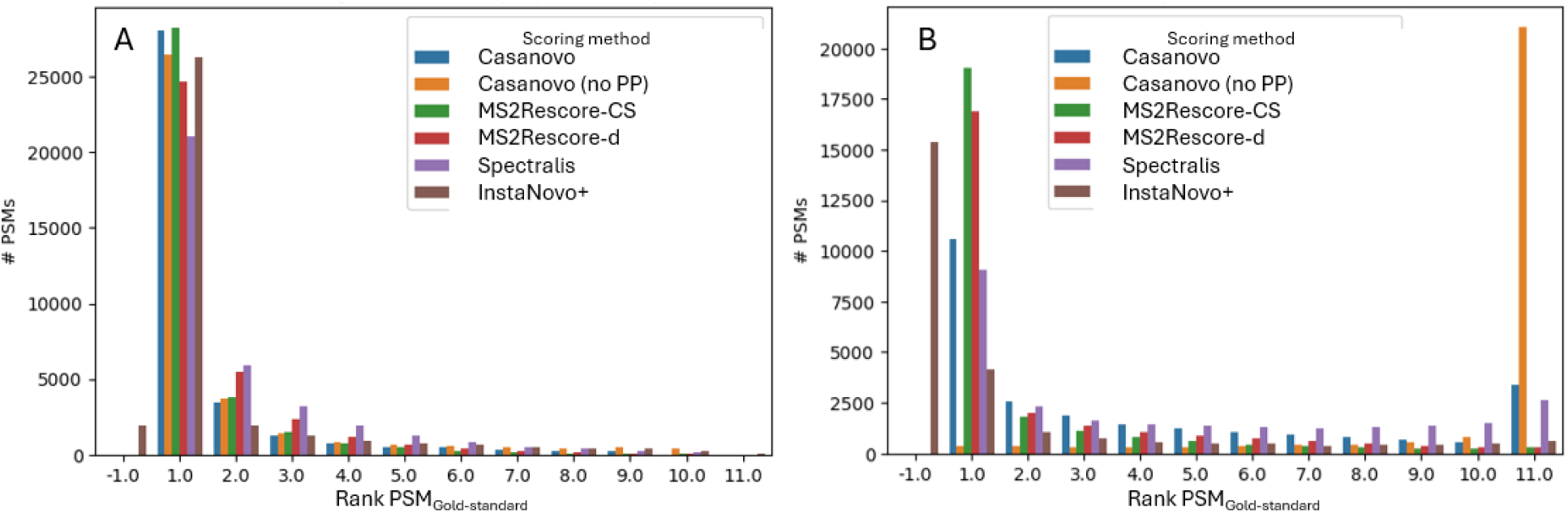
Summary of the rank of PSM_Gold-standard_ amongst the proposed candidates determined by several scoring methods for spectra where PSM_Gold-standard_ was (A) proposed in the top 10 candidates of Casanovo or (B) was absent. Scoring methods include Casanovo, Casanovo without precursor mass post processing (Casanovo no PP), MS2Rescore-CS, Ms2Rescore-d, Spectralis, and InstaNovo+.

These results bring two additional questions: (i) Why does Casanovo rank some candidates better than MS2Rescore-d, despite being outperformed in general, and (ii) why does Casanovo fail to include the correct PSM in its top ten candidates for 41.6% of spectra, whereas MS2Rescore-d can correctly flag the gold standard PSMs as the best match? These two topics are further investigated in the following sections.

To investigate why Casanovo is able to rank the gold standard PSM (*PSM_Gold-standard_*) better than MS2Rescore-d in *Spec_top10_*, the score distributions for gold standard and MS2Rescore-d top ranked PSMs (*PSM_MS2Rescore_*) are compared (Figure 8A, 8B). Casanovo scores for *PSM_Gold-standard_* are higher and follow a distinct distribution than *PSM_MS2Rescore_*, while the MS2Rescore-d scores between both PSMs are highly similar. This indicates that Casanovo is very certain the *PSM_Gold-standard_* is a better match, while MS2Rescore-d perceives both candidates as highly similar matches.

**Figure 8:**
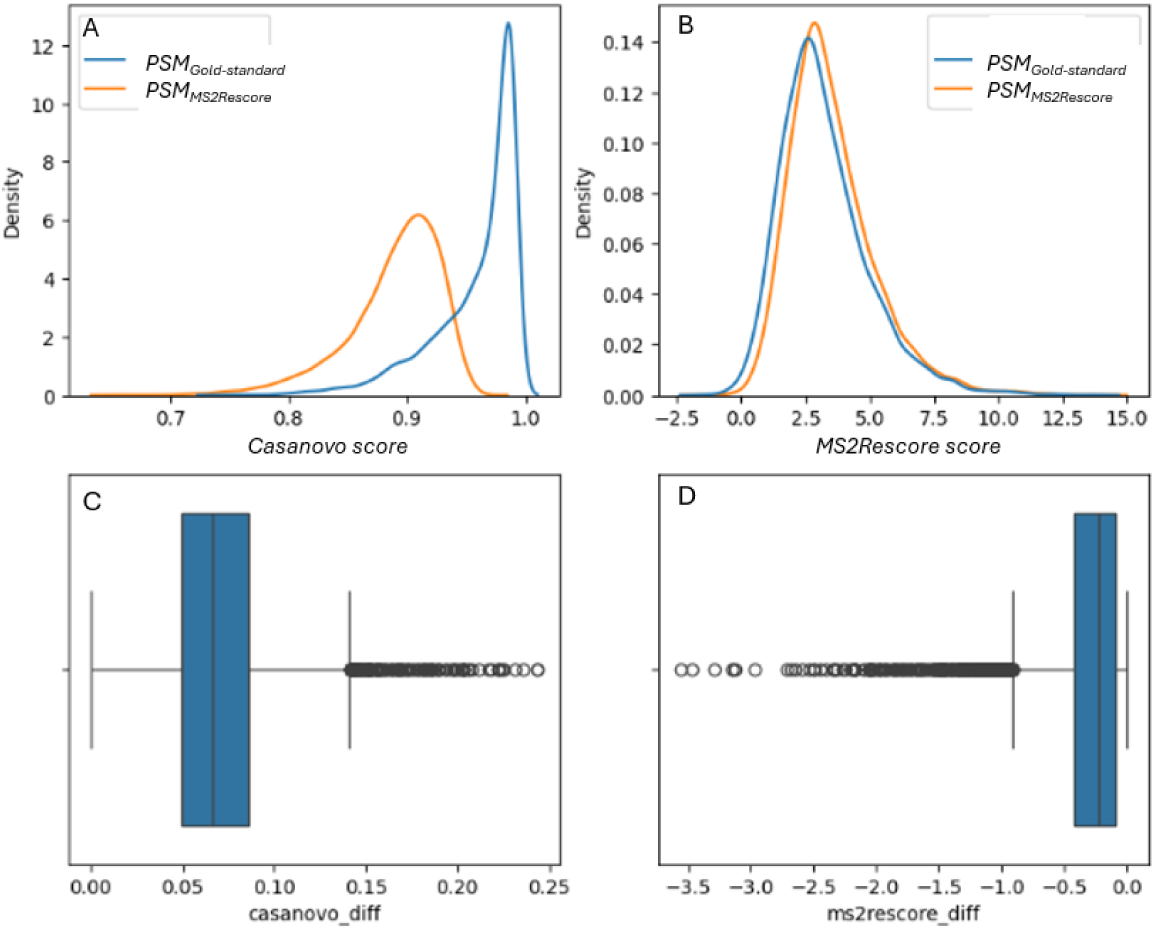
Score distributions of (A) Casanovo-scores, and (B) MS2Rescore-d scores for the (blue) PSM_Gold-standard_ and (orange) PSM_MS2Rescore_ in Spec_top10_. The score differences between the candidate PSMs (PSM_MS2Rescore_ - PSM_Gold-standard_) for (C) Casanovo-scores, and (D) MS2Rescore-d.

The Damerau-Levenshtein distance between both candidates indicates that most of these (92.0%) differ in at most three amino acid substitutions or transitions. 64.8% of these candidates are ambiguous, in the sense that the sites where the two peptide sequences differ are either both supported, or neither are supported, by fragment ion annotations in the spectrum. Moreover, 36.7% of the differences can be classified as a reordering of a small sequence of amino acids (e.g., VI vs VI, AI vs IA), 52.7% as isobaric variants (e.g., GG vs N, AG vs Q), and 10.5% as other differences, often mistakes such as isotopic errors, deamidations of amino acids, and close to isobaric variants. It is therefore surprising, as these errors are small and often not directly distinguishable based on fragment annotations, that Casanovo is still much more certain about *PSM_Gold-standard_* than *PSM_MS2Rescore_*.

While thus seemingly having higher discriminatory power between two highly similar candidates in *Spec_top10_*, Casanovo still fails to find the gold standard PSM in about half of database-identified spectra. To explain this discrepancy, spectral features between *Spec_absent_* and *Spec_top10_* were investigated as well. These features include the number of missing ion series, the hyperscore, the explained intensity, and the cosine similarity with the fragmentation spectrum predicted by MS2PIP. *Spec_absent_* show lower explained intensity (Figure 9A), more missing ion series (Figure 9B), slightly lower hyperscores (Figure 9C), and reduced cosine similarity to MS2PIP predictions (Figure 9D). Together, these indicators of noise and ambiguity likely prevent Casanovo’s beam-search from retrieving the correct PSM.

**Figure 9:**
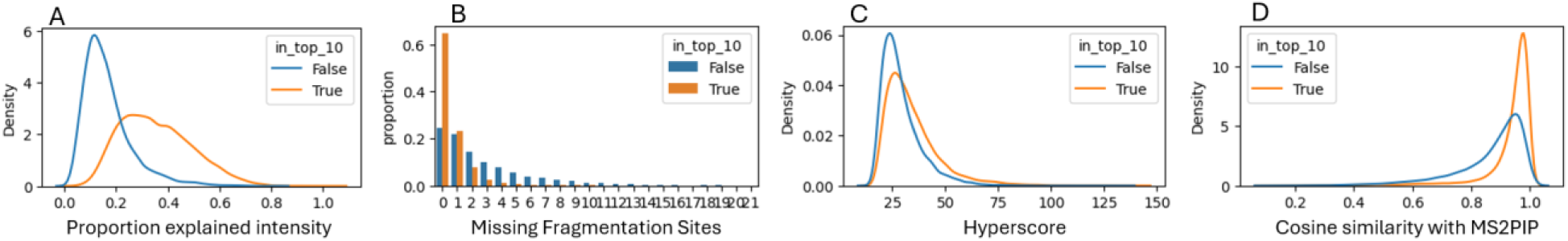
Feature distributions of PSM_Gold-standard_ for (blue) Spec_absent_ and (orange) Spec_top10_. (A) Ratio of explained intensity, (B) number of missing fragmentation sites, (C) hyperscore, and (D) cosine similarity with MS2PIP predictions.

Casanovo is not only unable to find *PSM_Gold-standard_* in *Spec_absent_,* it is also unable to detect *PSM_Gold-standard_* as the highest ranking candidate even when *PSM_Gold-standard_* is appended to the candidate list. To investigate why MS2Rescore-d is much more performant in this task, we leverage the interpretability of MS2Rescore by comparing the most important features used for training these models between the highest ranking Casanovo PSM (PSM_Casanovo_) and *PSM_MS2Rescore_* when MS2Rescore-d correctly ranked *PSM_Gold-standard_* highest. These features include the retention time error based on a comparison to predictions from DeepLC (Figure 10A), cosine similarity (Figure 10B), hyperscore (Figure 10C), and difference between the candidate and observed precursor mass in ppm (Figure 10D). From these feature differences, retention time error seemed to be the most important feature to discriminate these candidate PSMs. This reliance on the retention time feature indicates its importance for candidate selection, however, none of the *de novo* engines currently incorporate this as a feature.

**Figure 10:**
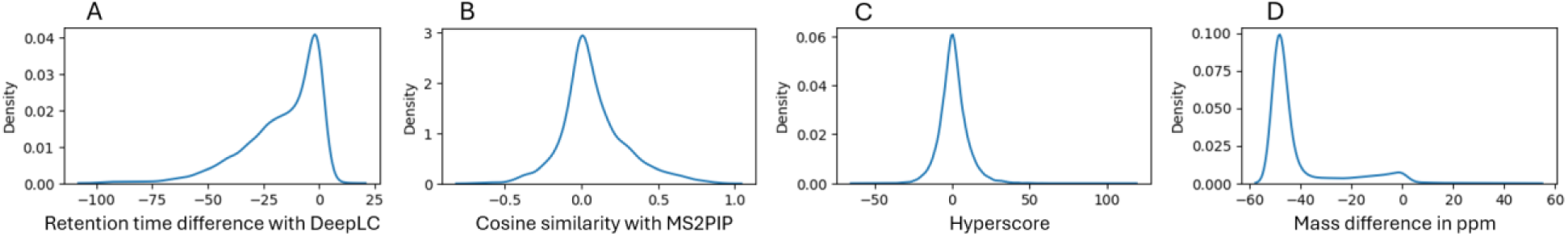
Feature differences between PSM_Casanovo_ and PSM_Gold-standard_ for spectra in Spec_absent_ when MS2Rescore-d scored PSM_Gold-standard_ highest. Positive values indicate that the feature value is higher in PSM_Gold-standard_. (A) Retention time difference with DeepLC predictions, (B) cosine similarity with MS2PIP predictions, (C) hyperscore, and (D) precursor mass difference in PPM (thresholded at +-50 ppm)

## 3 Error analysis of *de novo* PSMs

In previous sections, we assessed the accuracy of individual *de novo* models and the impact of post-processing strategies. Here, we shift focus to a detailed analysis of the errors that these models make. Specifically, we explore whether certain types of mistakes occur repeatedly, whether different models exhibit distinct error patterns, and whether these errors are limited to parts of the sequence that are unannotated by fragment ions. Finally, to estimate the upper bound of current *de novo* sequencing performance, we simulate an idealized ensemble by combining predictions from all eight models. By doing so, spectra which remain problematic for all models can be isolated, and will make clear which spectral features, such as noise and missing fragmentation sites, continue to limit *de novo* sequencing.

### 3.1 Sequencing errors

Across all eight *de novo* models, the distribution of peptide length-normalized Levenshtein distances between predicted and gold-standard sequences is bimodal: many predictions are either nearly correct or almost entirely different. To characterize the error types, the Levenshtein edit paths were parsed into mismatch regions (‘tags’; see Methods for details). Finally, the error types were grouped according to the nature of the tag pairs between the gold-standard and predicted sequences.

To gain an understandable overview of the errors, the tags were categorized in three sections: (i) single amino acid variants, (ii) localized errors with a maximum Damerau-Levenshtein distance of four, and (iii) large sequence mismatches with Damerau-Levenshtein distance larger than four. Both small and large tags were further categorized depending on the amino acid composition and isobaricity. An overview with the average proportion of error types across all models is shown in Figure 11. These categories are further elaborated below. Model-specific proportions for each error type are provided in the supplementary table T1 and will be referenced throughout the following sections.

**Figure 11:**
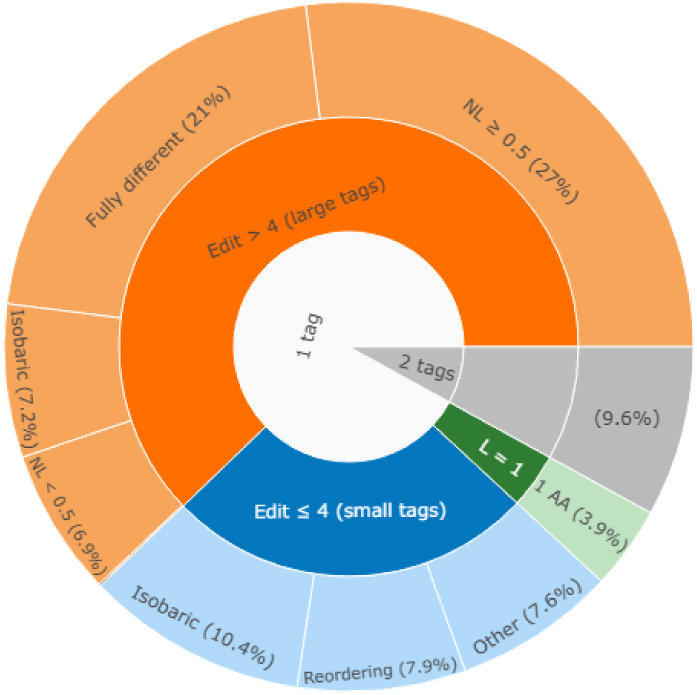
Overview figure of average occurrence of error types across all eight de novo models, subdivided into 3 layers of detail. Inner shell defines number of distinct mismatch regions found between the PSM_Gold-standard_ and PSM_denovo_ pair. Green, middle region indicates errors of single amino acid variants (Levenshtein distance = 1), blue regions indicates small mismatch regions with edit distance less than 5, further subdivided into amino acid permutations (reorderings), isobaric variants and non-isobaric variants. Orange regions indicate larger tags, subdivided into isobaric variants and three other groups depending on size of mismatch region (NL = peptide length-normalized Levenshtein distance).

### 3.2 Single amino acid variants

A minor section of errors consists of single amino acid variants (figure 12A). The percentage of errors belonging to this subsection varies widely across the models, from 1.48% for AdaNovo to 9.21% for π-PrimeNovo (Supplementary Table 1). The majority of these variants across all models consists of errors related to the deamidated forms of asparagine and glutamine. Interestingly, variants such as alanine to glutamine, and glycine to asparagine were amongst the most common errors made by, and unique to, PepNet (Figure 12D). These errors introduce a mass shift equal to a glycine (AG <-> Q, GG <-> N), highlighting a limitation of the non-autoregressive prediction approach of the model and associated post-processing algorithm to construct the peptide sequence from the amino acid probability matrix. Although only a single amino acid change could solve these errors, adding a single glycine would result in iterative changes in the selection of amino acids along PepNet’s probability matrix. Because these issues are easily resolved given the entire sequence, Spectralis and InstaNovo+ were able to correct this issue successfully as described in the previous section. It should be noted that, even though π-PrimeNovo also infers the peptide sequence from an amino acid probability matrix, it is better able to deal with this problem.

**Figure 12:**
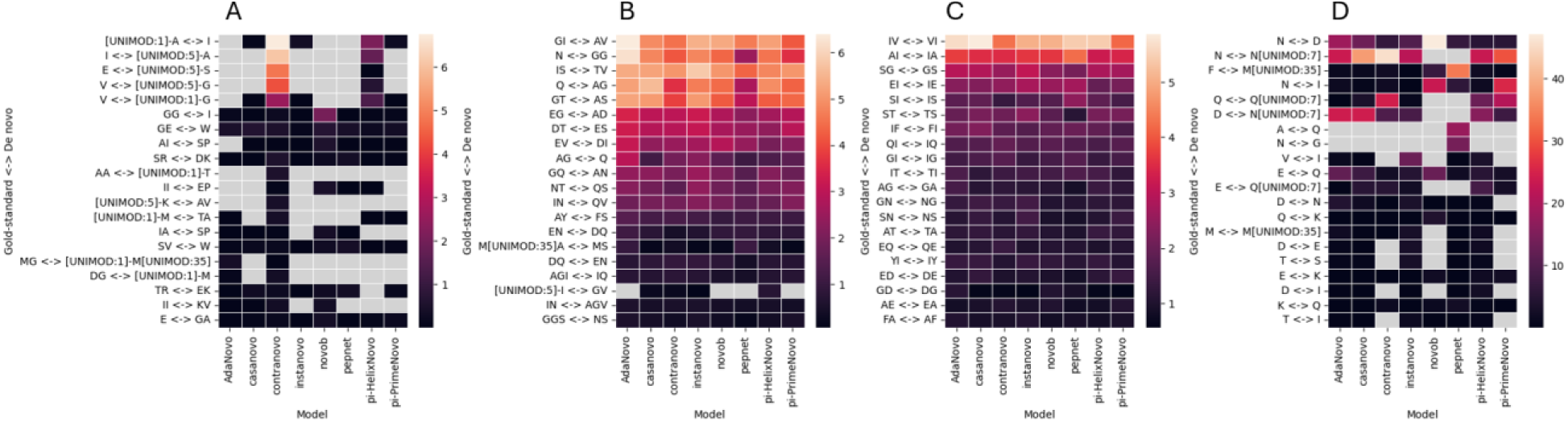
Most common tag pairs within each error type category colored by percentage of errors in each specific category. Error categories include (A) single amino acid variants, (B) amino acid permutations, (C) isobaric tags, and (D) the remaining tags.

**Figure 13:**
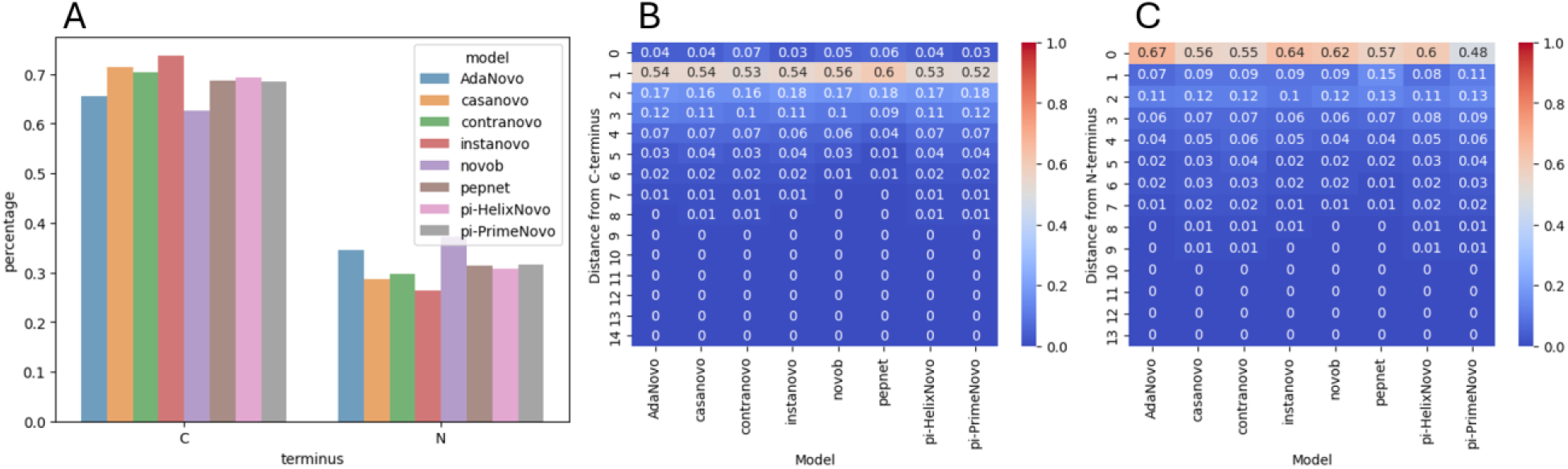
Location analysis for de novo PSMs with amino acid permutation and isobaric differences. (A) Ratio of tags residing either at the C-or N-terminus. (B) Starting site of the sequence mismatch, starting to count from the closest terminus either the C-or (C) N-terminus.

### 3.3 Localized errors

About 25% of errors are classified as single site, localized errors, defined by a Damerau-Levenshtein distance ≤ 4. These errors are further divided into three subcategories: (i) sequence permutations, i.e., tags with the same amino acid composition, (ii) isobaric tags, i.e., tags with a mass difference below 0.01 Dalton, and (iii) all other tags.

Sequence permutations occurred in, on average, 7.88% of all errors. The specific amino acids involved in permutation errors were largely consistent across models, with error frequency appearing to correlate with amino acid frequency rather than model-specific weaknesses (figure 12B). This error type is subtle, and the correct and incorrect sequences are quasi indistinguishable from each other when no fragment ions are present that indicate the correct positioning of these amino acids. Indeed, in the majority of cases (68.3%) with transpositions of two amino acids, no diagnostic fragment ions were present to distinguish between the two PSMs (Supplementary Figure S4a).

Localized isobaric errors were responsible for on average 10.4% of all errors. The most frequent isobaric differences are plotted in figure 12C. Interestingly, PepNet made significantly less AG to Q and GG to N errors. Similarly to sequence permutations, around half of the isobaric tags had no diagnostic ions to discern between the two cases (Supplementary Figure S4b).

Sequence permutations and isobaric tags are often indiscernible based on the fragment ion annotation. Next, we questioned where these tags mainly reside in the peptide sequence. To this end, the tags were attributed either to the amino (N) or carboxyl (C) terminus depending on which side the tag resides at, and the distance to this terminus was measured by counting the number of residues between terminus and tag. All models made these errors predominantly at the C-terminus (figure 13A). Furthermore, both at the N-and C-terminus, tags are mainly present at the terminus itself (minus the C-terminal lysine or arginine) (figure 13B, 13C). Importantly, most current *de novo* models start constructing the peptide sequence C-terminally, limiting the models to utilize sequence pattern information within this region and potentially negatively influencing the sequencing accuracy for the remainder of the sequence.

Lastly, in total 50,970 other variant pairs were found, encompassing on average 7.60% of all errors, with PepNet having vastly more of these errors (16.4%) than the other models. These tags are characterized by a mass difference larger than 0.01 Dalton. Even though in most models, PSMs with large precursor mass differences are filtered out, some models such as π-HelixNovo, Casanovo, AdaNovo, InstaNovo, and PepNet do still include them, resulting in the vast majority of these tags having a precursor mass difference larger than 1.1 Dalton. For the other models, the mass differences of these tags mostly include tags with mass differences below 0.1 Dalton. The remaining errors consist of tags with mass differences of around 1.0 Dalton, mimicking an isotopic error, and 0.984 Dalton, which corresponds to a deamidation. Note that these precursor mass differences also often occur in database searches.

Interestingly, most of the non-isobaric variants, except for PepNet and π-PrimeNovo, mainly occur at the N-terminus, in contrast with the isobaric variants (Supplementary Figure S5). This is unsurprising considering that a mass shift at the N-terminus does not change the m/z position of the y-ions, which are typically more prominent than b-ions. Nonetheless, non-autoregressive models do seem less biased towards this issue.

### 3.4 Extensive errors

The majority of errors encompass a large section of the peptide sequence, which we categorize here as a Damerau-Levenshtein distance larger than 4. We subdivided this error type in five sections: (i) amino acid permutations, (ii) isobaric tags, (iii) fully different peptide sequences, and non-isobaric tags encompassing (iv) less, or (v) more than half of the full peptide sequence.

In contrast to localized tags, these larger tags only show amino acid permutations in less than 0.1 % of cases. Instead, on average 7.23% of these errors consist of isobaric tags. NovoB (18.8%) and ContraNovo (12.7%) clearly had more of these error types.

On average, 21.0% of the wrong *de novo* predictions consisted of peptide sequences which were completely different from the ground-truth PSM. Although this could indicate the presence of a co-fragmenting precursor ion, the limited overlap amongst all tools as discussed in previous sections indicates there is little to no agreement to what the exact sequence of the co-fragmenting peptide would be.

## 4 The limits of current *de novo* models

To estimate the upper bound of current de novo sequencing capabilities, we simulated an idealized ensemble model: if any of the eight tools correctly predicts the peptide sequence for a given spectrum, the ensemble prediction is considered correct. While the best-performing individual model achieved an accuracy of 52.0%, the ensemble reached 67.9%. Furthermore, 10.3% of the incorrectly predicted peptides consisted of peptides present in the database, highlighting a potential alternative, better candidate.

To understand the factors limiting de novo performance, we examined several spectral and peptide properties. The most striking finding was the impact of missing complementary b-and y-ion fragments (Figure 14A). When all fragmentation sites were covered by at least one of these ions, 92% of the spectra were correctly predicted by the ensemble model. This dropped to 85%, 70%, and 50% when one, two, or three complementary ion pairs were missing, respectively. These results emphasize the reliance of current models on a complete fragmentation ladder. The influence of missing complementary ions was much more pronounced than the effect of peptide length, with longer peptides being more error-prone (figure 14B). Indeed, the effect of peptide length appears confounded by fragmentation completeness: as peptides get longer, the likelihood of missing complementary ions increases, making it difficult to disentangle the individual contributions of these two factors.

**Figure 14:**
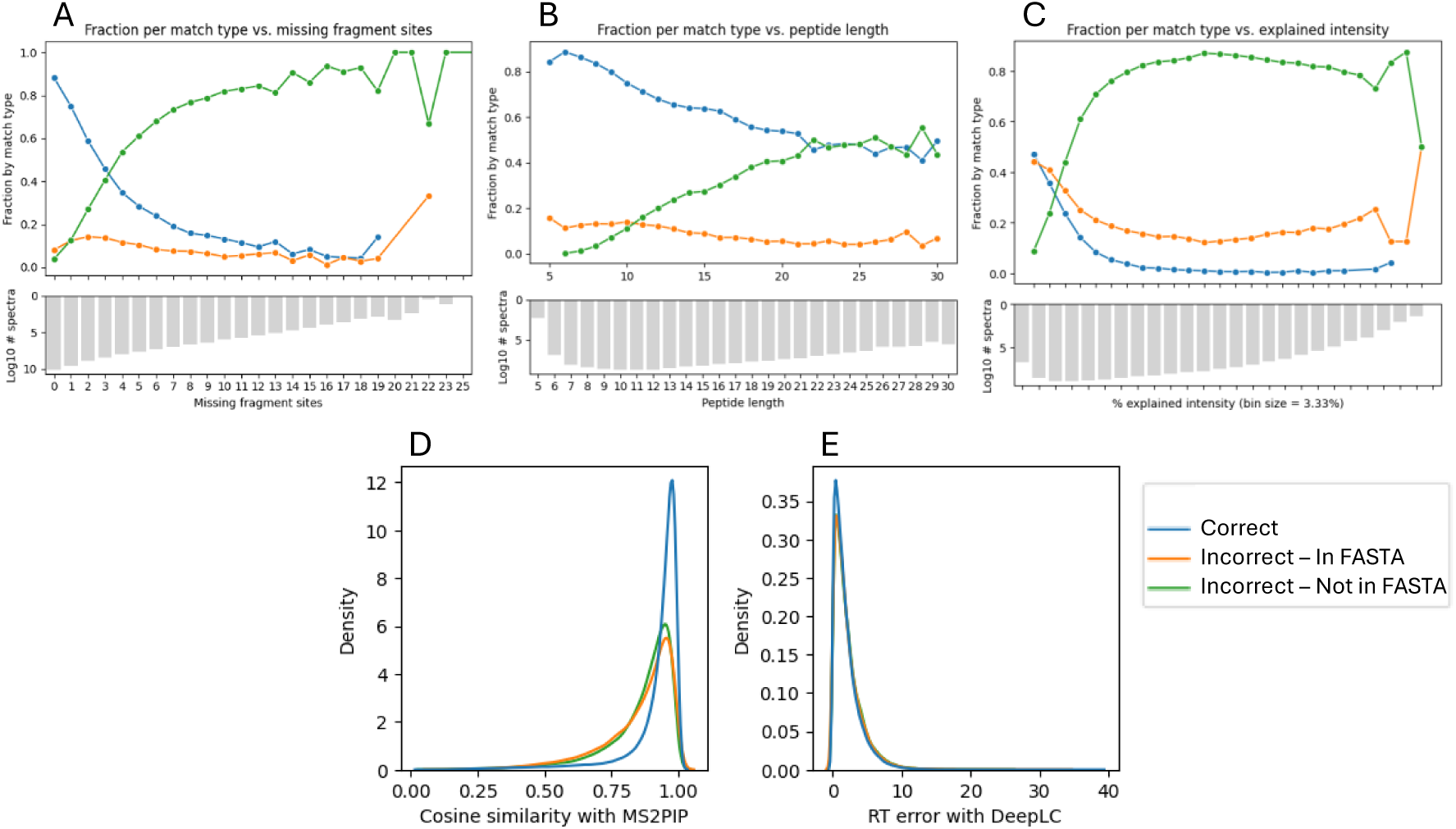
Spectral and peptide characteristics for spectra deemed (un)predictable by the ensemble model. Percentages for: (A) number of missing fragment ion pairs, (B) peptide length, and (C) bins of explained intensity. Feature distributions in relation to (D) MS2PIP, and (E) DeepLC predictions.

Spectral noise, quantified as the inverse of the percentage of explained intensity, was another strong predictor of failure. Only 17.0% of spectra with less than 5% explained intensity were correctly identified by any model, while this figure rose sharply to over 80% for spectra with more than 20% explained intensity (figure 14C).

To assess whether features orthogonal to MS2 fragmentation could help explain these limits, we used MS2PIP to compare predicted and observed intensity patterns (figure 14D), and DeepLC to assess retention time agreement (figure 14E). Spectra that all models failed to predict showed lower MS2PIP cosine similarity scores, suggesting these PSMs have less predictable or more noisy fragmentation patterns. Interestingly, although some PSMs suffer from sparse fragment ion ladders, low explained intensity, or erratic fragmentation, their retention-time deviations remain on par with those of well-predicted spectra. In other words, predictability of chromatographic behavior is largely independent of spectral quality.

These results highlight that current *de novo* tools struggle primarily when key fragmentation information is missing or masked by noise. Fragmentation intensity patterns help distinguish predictable from unpredictable spectra, while retention time offers little discriminatory power in this context. This sets a clear direction for future work: models must better handle incomplete fragmentation if possible, and should learn to exploit non-fragment-based features, such as retention time and ion mobility, if they are to surpass current limitations.

## Discussion

In this study, we benchmarked eight deep learning-based *de novo* sequencing models (AdaNovo^30^, Casanovo^29^, ContraNovo^31^, InstaNovo^32^, NovoB^33^, PepNet^34^, ϖ-HelixNovo^35^, and ϖ-PrimeNovo^36^), alongside three post-processing methods (Spectralis^37^, InstaNovo+^32^, and MS2Rescore^19^). Using common benchmarking metrics, peptide accuracy and recall, we evaluated their performance on a dataset acquired on QExactive and timsTOF instruments. While this dataset does not comprehensively represent all experimental conditions, our primary goal was to find a dataset that showed good performance for the *de novo* models, and use it to examine model behavior, error patterns, and their relationship to spectral features.

### Benchmarking landscape and limitations

In this benchmark, we focused on determining common limitations of current deep learning *de novo* models while developing a framework which makes execution and benchmarking the results of numerous models and refinement methods easy and reproducible. Unfortunately, as in previous efforts, our benchmark is not exhaustive, i.e., not all available models are included^45–48^. However, the way the benchmark is set up allows to easily include additional open source *de novo* models.

Compared to previous benchmarks, model rankings vary due to differences in datasets, metrics, and evaluation setups^28^. This highlights three key challenges: (i) the lack of consensus on benchmarking standards, (ii) the rapid development of new models, and (iii) the technical complexity of executing and comparing tools. Our NextFlow pipeline and Python API is easily extensible to other open source models and provides a robust way to execute models and to generate a wide range of evaluation metrics, addressing the second and third challenge effectively. Nonetheless, the extendibility is limited by the need for continuous maintenance. Additionally, the first two challenges require attention from the *de novo* community, as they require focused, domain-specific solutions.

To this end, we advocate for a centralized, community-driven benchmarking platform to support ongoing development and evaluation. ProteoBench^49^ is a promising candidate for this, as it offers infrastructure for public submission and transparent comparison, and uses clearly defined and community-accepted metrics. Such platforms can shift the focus from tool competitiveness to transparency and collaborative innovation. Potential risks, such as model overfitting to benchmark datasets, can be mitigated through community scrutiny and detailed reporting of model training and configuration. NovoBench^48^ is another notable example of a benchmarking platform, which additionally allows to retrain models when benchmarking, mitigating a limitation of ProteoBench.

### Performance and limitations of current models

Current de novo models perform remarkably well on HCD-acquired DDA data, with some exceeding 90% peptide-level accuracy on spectra with complete amino acid ladders. However, performance drops sharply on timsTOF data, due to the models having been trained on HCD-Orbitrap spectra. Similar issues likely affect ETD and other fragmentation methods^47^, highlighting a major limitation in model generalizability of currently pretrained models. While transfer learning on datasets with specific data properties may offer a cost-effective solution, it has not yet been widely explored in this context^22,50^.

Consistent with earlier studies, we observe that all models struggle with more ambiguous spectra due to lacking complementary ions or containing high noise^45–47,51^. These cases often involve multiple peptide candidates with highly similar scores, exacerbating the challenge of confident identification in a *de novo* context where the search space is maximized, and no database-based filtering can be applied to resolve ambiguities. Beyond modeling improvements, some research has explored the direct enhancement of data quality, for example, by using labeled fragmentation methods^52^, or by acquiring multiple spectra per precursor with different fragmentation methods^53^. Such acquisition strategies can help mitigate the current inability of computational approaches to confidently assign PSMs to ambiguous spectra^54^.

### Ambiguity, scoring, and confidence estimation

*De novo* sequencing hinges on ranking the correct peptide above all alternatives, a task that is directly influenced by the amount of fragment ion evidence. Furthermore, when considering only fragment ion evidence such as in classical PSM-scoring functions, for many spectra the correct PSM is not necessarily the highest scoring one^55^. Even in the context of database searching, dealing with this issue by dramatically reducing the search space is prone to compromising the FDR in the presence of ‘neighboring’ peptides, which are defined as irrelevant peptides with similar precursor mass and fragmentation spectra, as previously discussed^17,18,56^. A possible solution is thus to design better PSM-scoring functions that can succeed at assigning top scores to correct PSMs, even when less fragment ions are matched compared to alternatives. While some rescoring approaches attempt this by incorporating fragment intensity prediction^40^, Levenshtein distance estimation^37^, or reinterpret the mass spectrum using diffusion models^32^, their gains appear limited over state-of-the-art *de novo* models, likely because the *de novo* models already internalize these patterns during training.

Our results show that integrating orthogonal features, such as retention time, ion mobility, precursor delta mass, and peak matching statistics can enhance candidate discriminability. Ensemble approaches such as PostNovo also hold promise, recovering correct peptides missed by individual models^44^. This is further corroborated by our simulated, ideal ensemble that combines the strength of all eight models, which correctly predicted an additional 15.9% of all spectra. Nonetheless, in many cases the same issue still remains: the correct candidate needs to be discernible from the other high scoring candidates given the experimental data, which is not always feasible, no matter the accuracy of the PSM scoring function. Indeed, database search engines can only (partly) solve this issue by reducing the search space.

This sequence ambiguity problem is expected to become more pronounced as *de novo* engines begin to support PTM prediction. PTMs substantially increase the search space even more, making it more likely that multiple (modified) peptide candidates can explain the same spectrum equally well^57^. Without robust ways to estimate identification and modification localization confidence in an interpretive way, this added complexity risks inflating false positives, thus undermining trust in the predictions^58^.

In response to this challenge, several interpretability methods have been proposed to better understand model predictions. For instance, pi-xNovo visualizes attention matrices to highlight which spectral peaks most influence the predicted sequence^59^, while techniques like integrated gradients attempt to trace individual predictions back to distinct input features^60^. These approaches offer valuable insights into model behavior and can help identify whether decisions are driven by informative or spurious signals. However, such interpretability methods serve a fundamentally different purpose than statistical confidence estimation. While these enhance transparency and can guide model development, they do not provide a quantitative measure of how likely a prediction is to be correct. Indeed, these metrics lack the statistical rigor required to assess identification reliability for each spectrum, particularly in ambiguous or low-quality cases.

This distinction underscores the need for spectrum-level confidence estimation methods in *de novo* sequencing. In contrast to database searches, which benefit from the well-established target-decoy strategy, no widely accepted equivalent exists for de novo predictions. Instead, global PSM score thresholds are often applied to control precision, either arbitrarily or calibrated using ground-truth references^52^. Yet, we show that such thresholds can be misleading, as models may assign high, overconfident scores to incorrect predictions. Decoy spectrum-based approaches offer an interesting alternative to set adequate score-thresholds, but are also not spectrum-specific^47^.

To address this, it has been proposed to estimate spectrum-specific E-values, representing the expected number of peptides that would achieve a score equal to or better than the observed match by chance^55,61^. While this is conceptually similar to confidence estimation in database search tools like MS-GF+, applying such methods to *de novo* sequencing presents fundamental challenges. Most *de novo* models are optimized to return the highest scoring PSM. They do not evaluate or calibrate scores across the broader candidate space, nor are they optimized to separate the top prediction from other plausible alternatives. As a result, E-values calculated from such models can be artificially low, not because the model is confident in a statistically meaningful sense, but because it hasn’t assessed the uncertainty in the surrounding sequence space.

One potential path forward is to shift focus from the absolute score of the top prediction to the discriminative gap between top-scoring candidates, and based on this, assign a statistical measure as implemented in Johnson *et al*^62^. If the score difference between the best and the second-best sequence is large enough, we can be more confident in the assignment. Conversely, small differences may indicate ambiguity. This strategy could help filter results in a spectrum-specific way, particularly when combined with post-processing tools that resolve uncertain sequence regions by using orthogonal data such as a database^63,64^. In cases where multiple sequences share substantial overlap, it may be preferable to return a consensus sequence with variable regions annotated as uncertain, resembling tag-based approaches^3,65–67^.

### Toward practical applications

It is important to contextualize what current *de novo* sequencing can and cannot deliver. Only about half of the spectra identified by database search engines are also correctly identified by *de novo* methods. The rest either lack sufficient signal or represent cases with multiple indistinguishable peptide candidates.

In fields like metaproteomics or antibody sequencing, overconfidence in such PSMs from ambiguous spectra can distort downstream interpretation. However, existing strategies, such as alignment-based post-processing^8^ or taxonomic assignment using least common ancestor strategies^68^ are already equipped to handle partial or uncertain sequences^69^. The characterization of prediction errors by these models as was performed in this paper can further inform such approaches.

Thus, rather than expecting full-length, high-confidence *de novo* sequences for every spectrum, we argue for a more realistic goal: extracting reliable sequence tags or partial matches where possible, with explicit uncertainty annotation^66^. This allows downstream applications to leverage the information without being misled by overconfident assignments.

## Conclusion

In this study, eight *de novo* models and two refinement models were evaluated against database search results of well-characterized benchmark datasets as gold standard. Although model performance varied, a common subset of spectra were found to be trivial to predict accurately for each model. Moreover, we identified that the most informative feature for prediction errors was missing complementary ions. These errors mainly constitute of very large errors that span almost the entire peptide, or small, localized errors which are often isobaric and largely non-specific between the models. Although we show that orthogonal features such as retention time might be promising to help resolve such cases, future research in *de novo* sequencing should focus on strategies that can highlight sequence prediction uncertainty based on a measure of the inherent ambiguity in the fragmentation spectrum. Finally, we have implemented a computational pipeline that makes it easy to execute all of these models in a centralized fashion, and thus perform a similar analysis as presented here with any user-specified dataset, and/or additional *de novo* models as these become available.

## Materials and Methods

### Datasets

Raw files were downloaded from PXD028735, containing samples of lyophilized MassPrep *Escherichia coli* digest standard from Waters, Human K562, and Yeast protein digest extracts from Promega either analyzed separately or mixed in a 65% human, 15% yeast, and 20% E. coli ratios. Only the runs ran in DDA-mode and acquired on Thermo Orbitrap QE HF-X and Bruker timsTOF Pro instruments were used. MGF-files were downloaded from PXD023217 from the CAMPI-study. These samples consist of a mock-community of species and a human fecal microbiome. The samples were acquired on different instruments, including the Orbitrap QE HF, QE Plus, Fusion Lumos and a timsTOF and were all run in DDA mode. For more information about the sample preparation and data acquisition, see the original publications.

### Raw file processing

Orbitrap and timsTOF raw files from PXD028735 were converted to MGF format with MSConvert (version 3.0.24225-374f83c) and MGF files from PXD023217 were downloaded from PRIDE. Spectra with less than 15 peaks were filtered out and only the 800 most intense peaks in each spectrum were kept, as this is the maximum peaks the *de novo* models standardly use. To acquire the gold-standard peptide identifications for the spectra, the MGF files were searched with Sage and rescored with MS2Rescore (version 3.0.2). Cysteine carbamidomethylation was set as fixed, and methionine oxidation as variable modification. Also the following N-terminal modifications were set as variable: acetylation, carbamylation, ammonia-loss and the combination of carbamylation and ammonia-loss. For a comprehensive set of database search settings, we refer to the supplementary JSON-files. FASTA database for *Homo sapiens*, *Escherichia coli*, and *Saccharomyces cerevisiae* were downloaded from UniProt on the 31th of July 2024. *Homo sapiens* FASTA only includes reviewed and canonical proteins, while for *E. coli* and *S. cerevisiae* the pan proteomes were downloaded.

### *De novo* NextFlow pipeline

The *de novo* models included in this comparative analysis are AdaNovo, Casanovo, ContraNovo, InstaNovo(+), NovoB, PepNet, π-HelixNovo, π-PrimeNovo, and Spectralis. To streamline their execution and ensure reproducibility, we developed a modular pipeline in NextFlow that centralizes configuration and execution of all models. Pretrained models provided by the respective developers were used without retraining. Full model versions and search settings are provided on GitHub.

For downstream analysis, we created a Python API to load, benchmark, and visualize the de novo results in a uniform format. This framework is based on the psm-utils package^70^ and supports a hierarchical structure (Figure 15): PSMs with multiple score assignments are grouped per spectrum, and all spectra belonging to a raw file are stored in a Run object. Compatibility with psm-utils enables seamless evaluation of de novo results alongside database search outputs from a variety of search engines.

**Figure 15:**
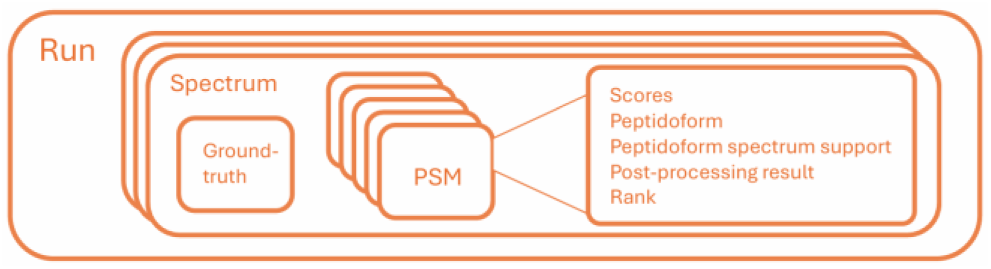
Schematic representation of the de novo results data object

### Post-processing *de novo* results

The pipeline includes three tools to post-process results of any of the supported *de novo* models. Spectralis and InstaNovo+ can rescore and refine candidate PSMs, using a Levenshtein distance estimator and a diffusion decoder respectively. MS2Rescore can rescore PSMs using matching features generated by machine learning models including DeepLC, MS2PIP, and IM2Deep as well as newly designed features which include an implementation of the hyperscore and statistics on missing complementary ion pairs.

The MS2Rescore framework was adapted to rescore *de novo* sequencing results and ideally requires database search results as well. In case database search results are available, these PSMs are rescored using MS2Rescore. A list of high-confident PSMs are used to calibrate and transfer learn the DeepLC and IM2Deep models. Both the calibration set and the models are stored and used for inferring retention time and collisional cross section predictions for *de novo* PSMs. After feature generation, Percolator models are trained on the database results and used for inferring new PSM-scores for *de novo* PSMs. In case no database search results are available, a selection of high confident *de novo* PSMs are selected as targets and lower-ranking PSMs are used as decoys, as similarly performed previously in approaches such as Nokoi^71^. Feature generator fine-tuning, Percolator model training and inference then proceeds the same way as described for database search results.

For reranking PSM candidates, a custom script was made so Casanovo can be run as a scoring function instead of a *de novo* model. Essentially, the model is run by inputting both the spectrum and a particular peptide sequence, returning both the peptide score and amino acid scores for this peptide.

### Evaluation metrics

To evaluate the performance of the *de novo* models, several evaluation metrics were used based on Bittremieux et al^28^ and originally proposed in Tran et al^72^. To evaluate a ‘match’ with the gold standard, the PSM assignments from the *de novo* model are compared with the database search results. A match is defined as a peptide which matches exactly with the ground-truth, while allowing leucine and isoleucine substitutions. If the *de novo* PSM does not match, yet predicted a peptide present in the FASTA database, it is annotated as ‘Incorrect – In FASTA’.

Precision is defined as in Bittremieux et al^28^: The number of true positives divided by all predictions. To generate the PC-curve, precision was calculated at different levels of coverage defined by setting thresholds on the peptide scores. Coverage is defined as the proportion of PSMs with scores above this score threshold amongst all predicted peptides or amongst all PSMs confidently identified with the database search engine at 1% FDR.

Levenshtein distance or edit distance was calculated by using the following packages: Levenshtein (https://github.com/rapidfuzz/Levenshtein) and nltk (https://tedboy.github.io/nlps/generated/generated/nltk.edit_distance.html).

MS2Rescore score differences were calculated by subtracting the MS2Rescore-scores of two PSMs generated by the pretrained Percolator models as described above.

### Tag generation for error analysis

Error tags were generated from the edit operations returned by the edit_distance function from nltk. Multiple tags within a peptide which are non-isobaric but become isobaric when merged, are merged. Additionally, the locations of the tags were determined by counting the number of amino acids from the closest terminus. Ambiguity regions between two PSMs are defined as tags which either lack all complementary ions in both PSMs or have at least a single fragment ion (b-or y-ion) at every fragmentation site in both PSMs.

### Simulated, ideal ensemble model

To evaluate the upper bound of current state-of-the-art de novo sequencing, we used a simulated ensemble model. This ensemble model takes the best candidate amongst the top candidates generated by any of the eight *de novo* models. For example, if only one model predicts the correct candidate, the ensemble model correctly predicted the spectrum.

## Code and data availability

Code and notebooks to generate the figures in the manuscript are available at https://github.com/SamvPy/DeNovo_Benchmark.

## Acknowledgments

TC, RB, and T.V.D.B. acknowledge funding from the Research Foundation Flanders (FWO) with grant numbers [12A8W25N, 12A6L24N, 1286824N]. LLE acknowledges support from the Research Council of Finland [341342, 364700], Sigrid Juselius Foundation, Cancer Foundation Finland, Biocenter Finland, and ELIXIR Finland. This work has benefited from collaborations facilitated by the Metaproteomics Initiative (https://metaproteomics.org/) whose goals are to promote, improve, and standardize metaproteomics^73^.

