## Supplementary Figures for "Limitations of *de novo* sequencing in resolving sequence ambiguity"

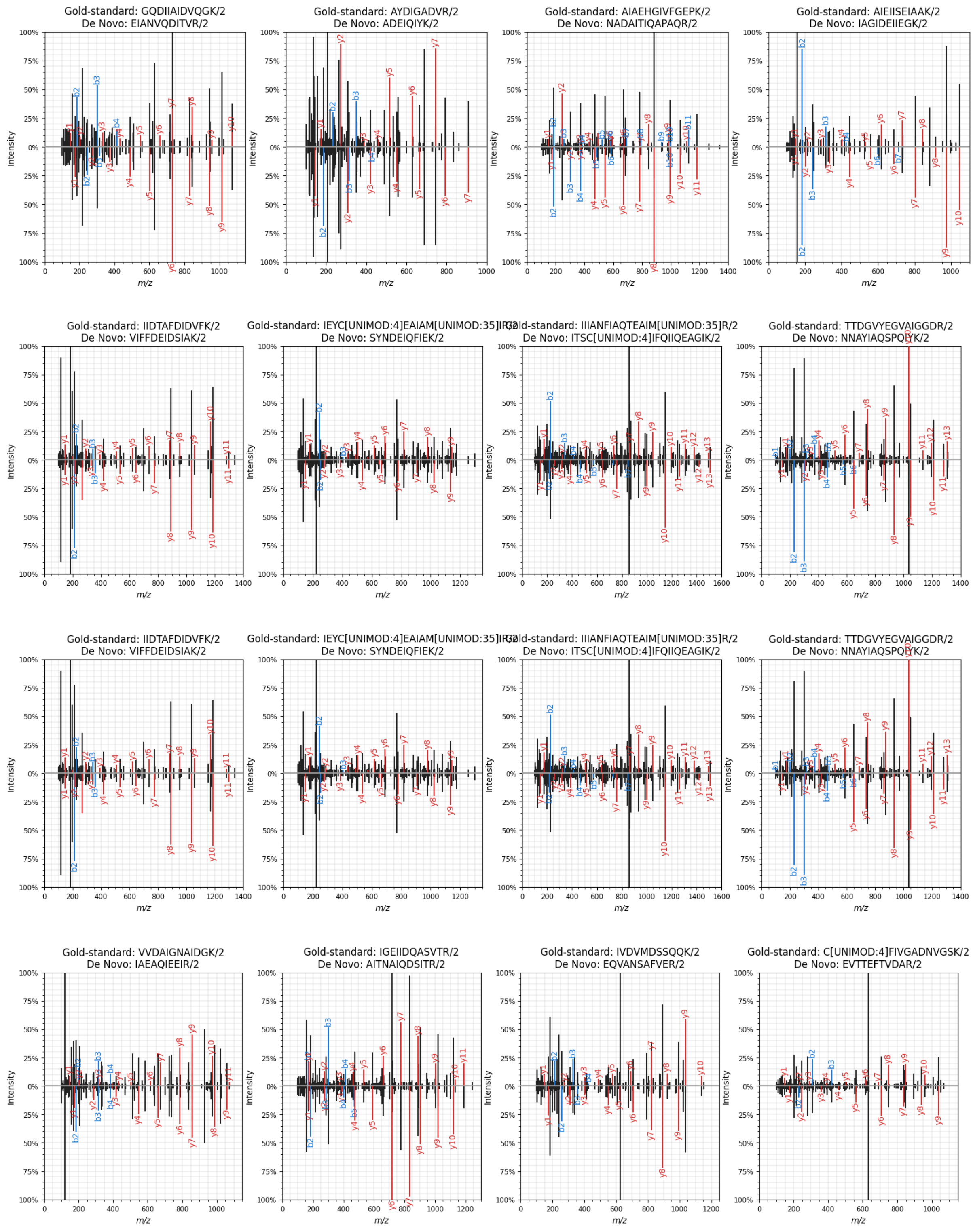

Gold-standard: IIDTAFDIDVFK/2
De Novo: VIFFDEIDSIAK/2
Gold-standard: IEYC[UNIMOD:4]EAIAM[UNIMOD:35]IR/2
De Novo: SYNDEIQFIEK/2
Gold-standard: IIIANFIAQTEAIM[UNIMOD:35]R/2
De Novo: ITSC[UNIMOD:4]IFQIIQEAGIK/2
Gold-standard: TTDGVYEGVAIGGDR/2
De Novo: NNAYIAQSPQYK/2


*S1: Annotated mirror plot of spectra which acquired de novo predictions, where all* de novo *models predict the same peptide sequence.*

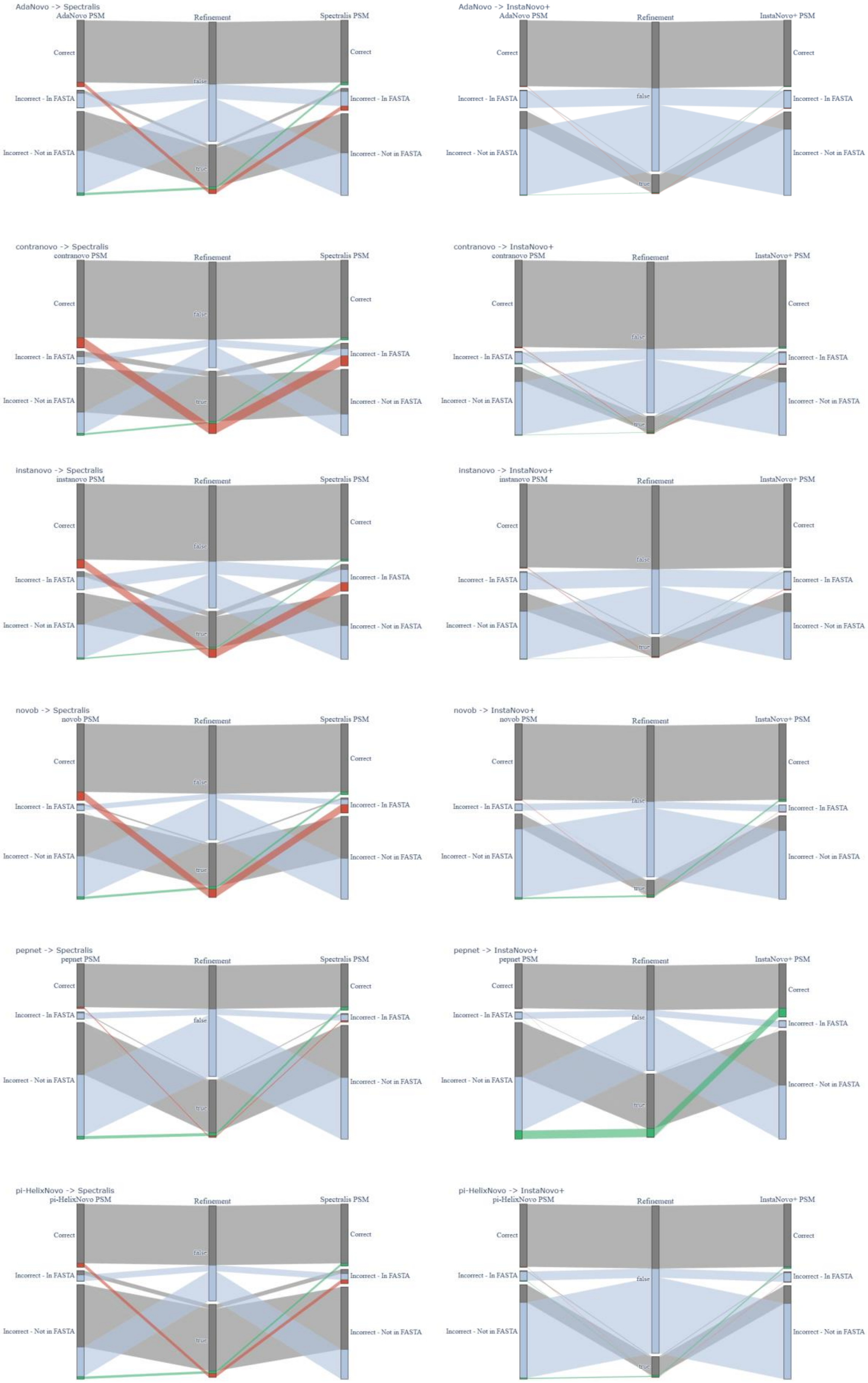
AdaNovo -> Spectralis
AdaNovo PSM
Refinement
Spectralis PSM
Correct
Incorrect - In FASTA
Incorrect - Not in FASTA
false
true
AdaNovo -> InstaNovo+
AdaNovo PSM
Refinement
InstaNovo+ PSM
contranovo -> Spectralis
contranovo PSM
contranovo -> InstaNovo+
instanovo -> Spectralis
instanovo PSM
instanovo -> InstaNovo+
novob -> Spectralis
novob PSM
novob -> InstaNovo+
pepnet -> Spectralis
pepnet PSM
pepnet -> InstaNovo+
pi-HelixNovo -> Spectralis
pi-HelixNovo PSM
pi-HelixNovo -> InstaNovo+

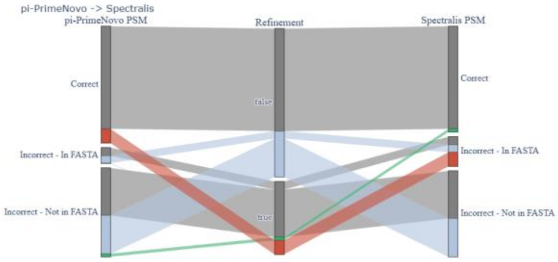


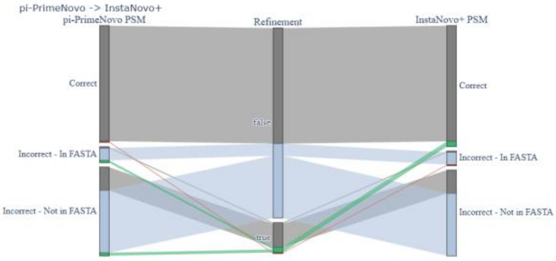


*S2: Sankey plots showing the effect of refinement for (left) Spectralis and (right) InstaNovo+ on PSM-correctness for* de novo *PSMs generated by (A) AdaNovo, (B) ContraNovo, (C) InstaNovo, (D) NovoB, (E) PepNet, (F) π-HelixNovo, and (G) π-PrimeNovo*

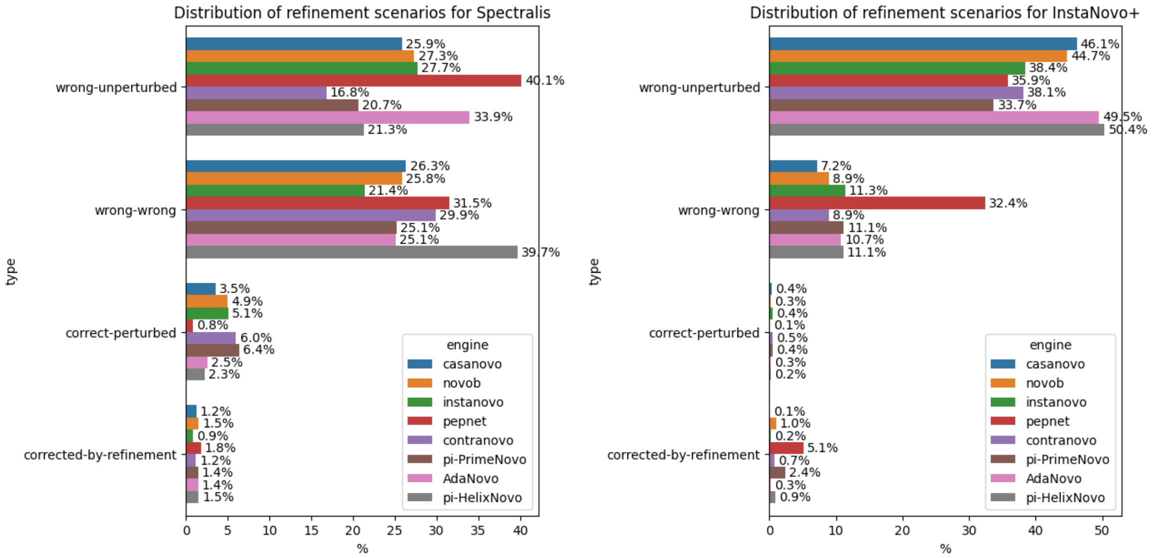


*S3: Detailed plot of Figure 5b. Amount of refinements falling in given refinement scenario for each* de novo *model separately after using Spectralis (a) or Instanovo+ (b).*

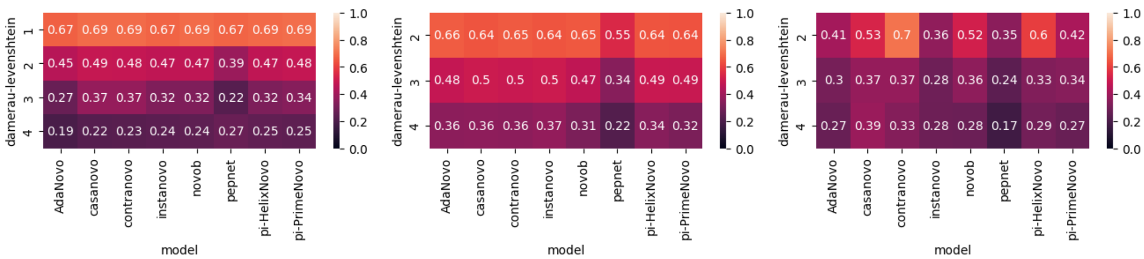


*S4: Ambiguity map (indicating proportion of tags defined by missing all complementary ions are having at least one complementary ion spanning the entire tag) for localized, (a) permutation tags, (b) isobaric tags, and (c) non-isobaric tags.*

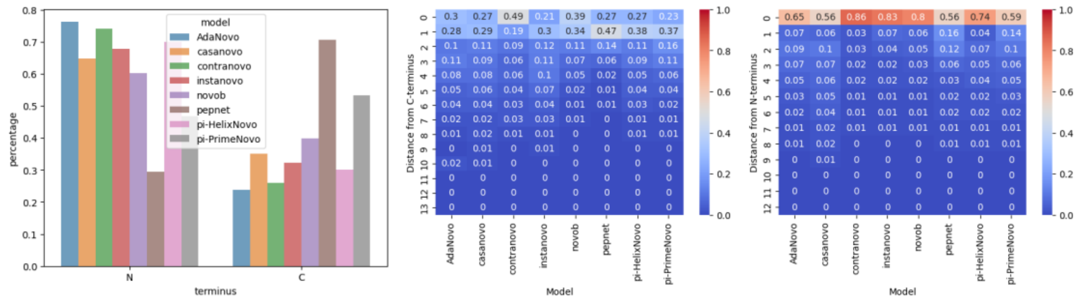


*S5: Location analysis for de novo PSMs with amino acid permutation and isobaric differences. (A) Ratio of tags residing either at the C- or N-terminus. Starting site of the sequence mismatch, starting to count from the closest terminus either the (B) C- or (C) N-terminus.*

### Supplementary tables

*T1: Occurrence of localized error types for each model amongst four categories as a percentage of all errors made by each model.*

| Model name | SAAV (%) | Permutations (%) | Isobaric variants (%) | Non-isobaric variants (%) |
|---|---|---|---|---|
| AdaNovo | 1.48 | 7.30 | 9.30 | 4.68 |
| Casanovo | 1.81 | 8.86 | 11.8 | 3.64 |
| ContraNovo | 4.21 | 9.40 | 11.9 | 6.09 |
| InstaNovo | 2.34 | 7.79 | 9.66 | 7.10 |
| NovoB | 2.02 | 9.30 | 12.4 | 6.87 |
| PepNet | 6.56 | 3.85 | 6.19 | 16.4 |
| π-HelixNovo | 3.83 | 6.27 | 9.08 | 7.14 |
| π-PrimeNovo | 9.21 | 10.2 | 12.5 | 8.88 |
| AVERAGE | **3.93** | **7.87** | **10.4** | **7.60** |

*T2: Occurrence of large error types for each model amongst four categories as a percentage of all errors made by each model.*

| Model name | Permutations (%) | Isobaric variants (%) | Fully different (%) | DL < half peptide length (%) | DL > half peptide length (%) |
|---|---|---|---|---|---|
| AdaNovo | 0.07 | 3.11 | 25.1 | 6.72 | 31.7 |
| Casanovo | 0.08 | 4.57 | 23.6 | 5.37 | 30.3 |
| ContraNovo | 0.11 | 12.7 | 24.6 | 3.84 | 21.8 |
| InstaNovo | 0.06 | 2.70 | 23.0 | 6.90 | 31.0 |
| NovoB | 0.14 | 18.8 | 16.7 | 5.58 | 20.2 |
| PepNet | 0.03 | 3.41 | 17.5 | 10.9 | 30.5 |
| π-HelixNovo | 0.07 | 5.27 | 22.1 | 6.81 | 29.4 |
| π-PrimeNovo | 0.13 | 7.36 | 15.8 | 9.17 | 21.3 |
| AVERAGE | 0.09 | 7.23 | 21.0 | 6.91 | 27.0 |
